# The α/β hydrolase domain-containing protein 1 (ABHD1) acts as a lysolipid lipase and is involved in lipid droplet formation

**DOI:** 10.1101/2023.12.17.572040

**Authors:** Ismael Torres-Romero, Bertrand Légeret, Marie Huleux, Damien Sorigue, Alicia Damm, Stéphan Cuiné, Florian Veillet, Carla Blot, Sabine Brugière, Yohann Couté, Matthew G. Garneau, Hari Kiran Kotapati, Yi Xin, Jian Xu, Philip D. Bates, Abdou Rachid Thiam, Fred Beisson, Yonghua Li-Beisson

## Abstract

Lipid droplets (LDs) are the major sites of lipid and energy homeostasis. However, few LD biogenesis proteins have been identified. Here, using *Chlamydomonas* as a model, we show that ABHD1, a member of the α/β hydrolase domain-containing protein family, is a novel type of LD-associated protein which stimulates LD formation through two distinct actions on the LD surface, one enzymatic and the other structural. ABHD1 was localized to LD surface in *Chlamydomonas* cells. The knockout mutants contained similar amounts of triacylglycerols (TAG) but their LDs showed an increased content in lyso- derivatives of the betaine lipid diacylglyceryl-*N,N,N*-trimethylhomoserine (DGTS). Over-expression of *ABHD1* in Chlamydomonas induced LD formation and boosted TAG content, suggesting a key role in LD biogenesis. The purified recombinant ABHD1 protein hydrolyzed lyso-DGTS, producing a free fatty acid and a glyceryltrimethylhomoserine moiety. In vitro experiments using droplet-embedded vesicles showed that ABHD1 promoted LD emergence. Taken together, these results identify ABHD1 as a new player in LD formation by its lipase activity on lyso-DGTS and by its distinct biophysical property. This study further suggests that lipases targeted to LDs and able to act on their polar lipid coat may be interesting tools to promote LD assembly in eukaryotic cells.

**Significant statement:** Lipid droplets are subcellular organelles specialized for triacylglycerol storage. Their dynamic turnover is key to managing energy homeostasis in response to cell cycle states and environmental cues. To gain insights into LD biogenesis, we characterized a putative α/β- hydrolase (ABHD1) in the model algae *Chlamydomonas reinhardtii* and show it is located at the LD surface. We found that ABHD1 overexpression promotes LD formation and acts as a lipase mainly on lyso derivatives of the betaine lipid diacylglyceryl-*N,N,N*-trimethylhomoserine (DGTS), the major lipid constituent of the LD hemi-membrane. We also show that ABHD1 has a remarkable biophysical property favoring LD budding. This work thus identifies a novel type of lipase acting on betaine lipid and provides a first example of a protein with a dual function nvolved in LD formation.

## Introduction

Lipid droplets (LDs) are the site of lipid storage accumulation in eukaryotic cells. LDs are made of a neutral lipid core, consisting of mostly triacylglycerols (TAGs), and are surrounded by a monolayer of polar lipids (hemi-membrane) embedded with proteins (1). LDs evolved to sequester the non-membrane forming neutral lipids i.e. triacylglycerol (TAG) away from membranes. In addition to its major role as carbon storage, LDs contribute to maintain energy homeostasis, prevent lipotoxicity, act as a temporary depot for acyl chains and participate in lipid signaling (2–5). Thermodynamically speaking, the oil-in-water configuration of LDs sets them apart from all other subcellular organelles. Their formation therefore requires a tight coordination between the biophysical properties of their neutral lipid core, amphipathic polar lipids, and those of their protein coat. Accumulating evidence points to the importance of both proteins and lipids in maintaining a thermodynamically stable LD configuration and population (6–8).

LD biogenesis occurs in response to environmental cues (such as light, nutrition, temperature) or is developmentally regulated (as in seeds or in aging cells). Considering the paramount importance of TAG metabolism for human health and for biotechnology, LD formation has been studied intensively in various organisms ranging from yeast to mammals, land plants, and microalgae (3, 9). Current understanding is that the LD, at least most of it, emerges from specific subdomains of the endoplasmic reticulum (ER) membrane defined by the enrichment of various proteins including the protein SEIPIN found in mammals, plants, and fungi (10). Initial membrane lipid remodeling and TAG synthesis results in the formation of an “oil lens” between the leaflets of the ER bilayer which is then followed by its growth, and finally it pinches off the membrane toward the cytosol giving rise to a LD (6). It is worth pointing out that new evidence suggests that LDs may not necessarily ever pinch off from the ER, but rather remain attached to the ER (11). Up to now, only a handful of proteins have been demonstrated to play a role in LD biogenesis in plants, animals or yeast. These include the PERILIPIN1, FSP27 (Fat-specific protein 27), FIT2 (Fat storage-inducing transmembrane protein 2), the FATP1-DGAT2 (Fatty acid transport protein 1-diacylglycerol acyltransferase 2) complex, the LD-associated protein (LDAP), and finally SEIPIN with its interactors, i.e., LD-assembly factor 1 (LDAF), the LDAP-interacting protein (LDIP), or the membrane-tethering protein VAP27-1 (vesicle-associated protein 27-1) (9, 12–20).

In the past 10 years, *Chlamydomonas reinhardtii* has been used as a model green microalga to interrogate LD biogenesis and turnover (2). LD formation and degradation in *Chlamydomonas* can be manipulated easily, for example, by simply changing the nitrogen (N) level in the media (21). N starvation is one of the most potent triggers in inducing TAG accumulation, which results in a drastic increase in both LD number and size in algal cells (22). In addition to research on environmental conditions or genetic basis of TAG metabolism and regulation (12, 21–24, 24–32), the protein and lipid composition of the LD coat has also been reported (30, 33–35). The monolayer lipid-coat of a LD is made predominantly of the betaine lipid diacylglyceryl-*N,N,N*-trimethylhomo-Ser (DGTS), followed by sulfoquinovosyl diacylglycerol (SQDG) and phosphatidylethanolamine (PE), and small amounts of galactolipids (30). Over 200 proteins were found in a LD-enriched fraction in Chlamydomonas (36), but only a few have been characterized. This includes: the major lipid droplet protein (MLDP), considered as a structural protein of Chlamydomonas LDs (34, 37), the betaine lipid synthase BTA1 (38), the long-chain acyl activating enzyme LCS2 (39, 40), and various lipid trafficking proteins (41). No protein involved in LD budding and expansion has been identified yet in algae. To investigate LD biogenesis in Chlamydomonas, we have focused on a putative α/β hydrolase domain-containing protein (ABHD1), which is present in the proteomics of LDs isolated from N-starved and high light-exposed cells (30, 33). Using cell biological, biophysical as well as genetic (gain- and loss-of-function) and biochemical studies in Chlamydomonas, we demonstrate here that ABHD1 is a lipase mostly acting on lyso-DGTS that can promote LD formation in vivo and in vitro. A tentative model describing the dual role- enzymatic and non-enzymatic- of ABHD1 in the mechanism of LD formation is also proposed.

## Results

### Gene expression, structural and evolutionary features of ABHD1 protein

Searching public databases on mRNA expression levels for *ABHD1*, one can observe that its gene expression is increased twofold upon N starvation (42). Furthermore, our immunoblot using anti-ABHD1 antibodies showed that ABHD1 significantly increased in Chlamydomonas cells during N deprivation, a condition promoting TAG synthesis and LD formation (**Fig. 1*A***). Taken together these data strongly suggest that ABHD1 may be involved in LD biogenesis in Chlamydomonas. In addition to ABHD1, Chlamydomonas genome encodes another protein (Cre01.g010550) showing 51% identity to ABHD1, and we named here as ABHD2. ABHD2 is absent in the previously published LD proteomics for Chlamydomonas, and moreover its expression is not regulated by N deprivation (33, 42), therefore ruling out a possible complementary role in terms of LD biology.

**Figure 1.**
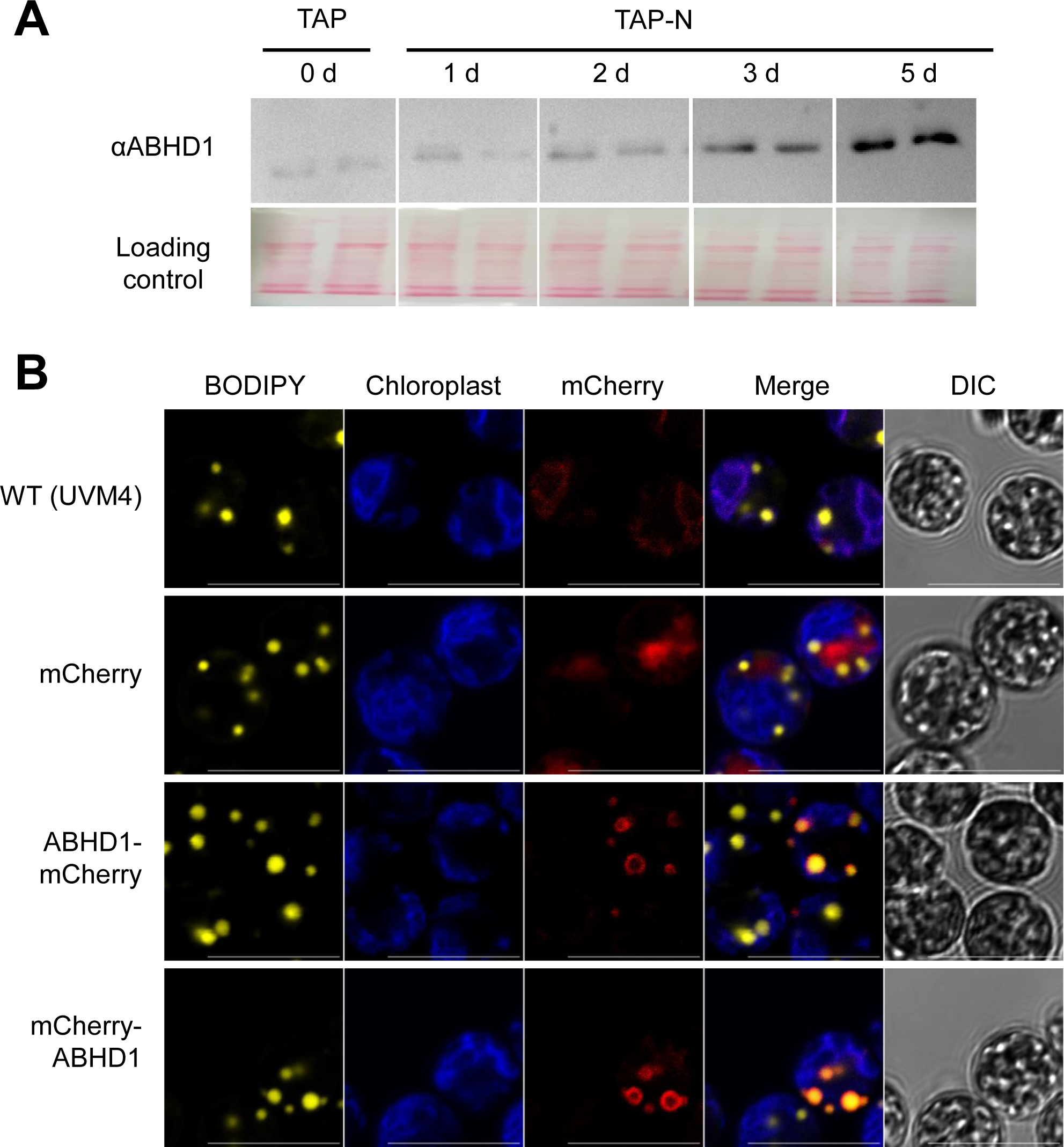
ABHD1 protein is located on the surface of lipid droplets and its quantity increases with prolonged N starvation. **A.** ABHD1 protein level as detected by immunoblot analysis using anti-ABHD1 antibodies. Total protein extracts (from 0.4 million cells) were loaded onto each lane. Ponceau’s stained proteins transferred onto nitrocellulose membrane were shown as control for protein load (whole lane displayed shrunk). Abbreviations: α, antibody; d, day; N, nitrogen; TAP, tris-acetate-phosphate media. **B.** Confocal imaging. Cells were cultured in TAP-N (1 d) and stained with BODIIPY to observe LDs. These are representative images from observations of >1000 cells from 10 independent transformation events. Pseudo-colors were used: BODIPY-stained LDs in yellow, chlorophyll autofluorescence in blue, mCherry and mCherry-tagged ABHD1 in red. DIC: differential interference contrast. Bar = 10 µm.

To gain insight on ABHD1 protein functions, we examined its primary amino acid sequence, its hydrophobicity (43) and its predicted structural features using alpha-fold (44). ABHD1 is composed of three major parts: an N-terminal transmembrane domain, one central α/β-hydrolase fold-containing domain, and an intrinsically disordered domain at its C-terminus (***SI Appendix*, Fig. S1*A-C***). In addition, the occurrence of a hydrophobic helix (alanine-rich domain) at its C-terminus (AA 396-490) (***SI Appendix*, Fig. S1*B*, *C***) could possibly serve as an anchor to LDs. Similar structural motifs at the C-terminus of the phosphatidylserine-specific phospholipase A1 have been shown to bind lipids (45). In terms of hydrophobicity (***SI Appendix*, Fig. S1*E***), ABHD1 shares similar overall hydrophobic features to MLDP but is a bit less hydrophobic (Gravy index of 0.05 comparing to 0.11 for MLDP) and does not contain the major hairpin hydrophobic domain present in in oleosins (34).

A phylogenetic tree based on amino acid sequence was constructed to explore the evolutionary history of ABHD1, which included 92 representative proteins from different clades of life (***SI Appendix*, Table S1**). The closest homologs to ABHD1 grouped in a clade distinct from bacterial and cyanobacterial homologues and this clade only included algae of the Chlorophytina taxon, which is composed of the classes Ulvophyceae, Trebouxiophyceae and Chlorophyceae (***SI Appendix*, Fig. S2**). ABHD1 homologs are absent in plant or mammalian cells, therefore ABHD1 is specific to Chlorophyta. The lack of homology in these different lineages seems to have been a rule rather than an exception for LD-associated proteins such as, for example, the protein Delayed in TAG Hydrolysis 1 (DTH1) (32). Increasing evidence seems to point out that organisms from distinct lineages equip their LDs with unique sets of proteins that do not share primary sequence homology but follow the overarching principles in LD structure/architecture and function.

### ABHD1 coats the entire LD and its over-expression induces LD formation

To investigate the functional role of ABHD1, we first sought to confirm that the protein was actually associated to LDs in vivo and determine whether it is its major or only location. We therefore fused the fluorescent protein mCherry at the N- or C-terminus of the ABHD1 protein (***SI Appendix*, Fig. S3*A*; Table S2**) and transformed the nuclear genome of Chlamydomonas with a construct expressing this chimeric protein under the strong constitutive PSAD promoter. To facilitate LD identification under the confocal microscope, N-deprived cells were examined and stained with the lipid dye BODIPY. More than 500 antibiotic resistant transformants (Both N and C terminal fusions) were screened based on mCherry signal. The mCherry signal was detected all around LDs, regardless of the N- or C-terminal position of the tag (**Fig. 1*B*; *SI Appendix*, Fig. S4 and S5, Movie S1**). Nevertheless, a higher number of positive lines were obtained when mCherry was fused to the N-terminus of ABHD1 protein. The fusion of mCherry to the C-terminus of ABHD1 seems to be less stable, leading to a higher rate of truncated or misfolded chimeric protein. If the disordered C-terminus region is important for proper folding and 3D structure or targeting to LD, this could explain the fewer amounts of over-expressors identified (***SI Appendix*, Fig. S1*B***). We have therefore focused on the lines where mCherry is fused to N-terminus of the ABHD1 protein to minimize misfolded proteins, such as aggregates or mis-targeted protein, that could occasionally be observed via mCherry signal in the cytosol. A more likely reason for mCherry signal in the cytosol is the use of the strong PSAD promoter, which resulted in potential ABHD1 over-expression. The robust signal emitted by mCherry fluorescence around the LD surface strongly indicates that ABHD1 is essentially a LD-associated protein and coats the entire LD (**Movie S1**).

While performing the subcellular localization experiment, we noticed LDs seem to be more abundant in *ABHD1* overexpressing lines during optimal growth than in WT cells (**Fig. 2*A***). Lipid analysis revealed that, while there was no difference in TAG content during N starvation between the *ABHD1* overexpressors (OEs) and the parental strain UVM4 (***SI Appendix*, Fig. S3*B***), OEs contained more TAGs during N replete growth (**Fig. 2*B***). Interestingly, the number of LD per cell was also altered, with a shift toward a higher number of LDs per cell in the OEs compared to the parental line (**Fig. 2*C***). These results thus indicate that in *C. reinhardtii* ABHD1 is not a limiting factor for LD accumulation under N-depletion but can induce the formation of LDs under N-replete conditions when overexpressed. In order to see if ABDH1 could induce LD formation in other cells than a microalga, we also expressed *ABHD1* in the quadruple yeast mutant H1246 strain where TAG synthesis is greatly reduced. *C. reinhardtii* ABHD1 (CrABHD1) expression alone in the *Saccharomyces cerevisiae* H1246 mutant resulted in a significant increase in TAG compared to the vector control line (**Fig. 2*D***). This increase in TAG was not as dramatic as with the native *S. cerevisiae* TAG-synthesizing enzyme DiacylGlycerol Acyltransferase 1 (ScDGA1), but it was nevertheless not negligible. This result suggests that the mechanism of LD induction on which ABDH1 acts exists in yeast. It also rules out that the observed TAG increase in Chlamydomonas ABDH1 OEs is due to the presence of the mCherry tag.

**Figure 2.**
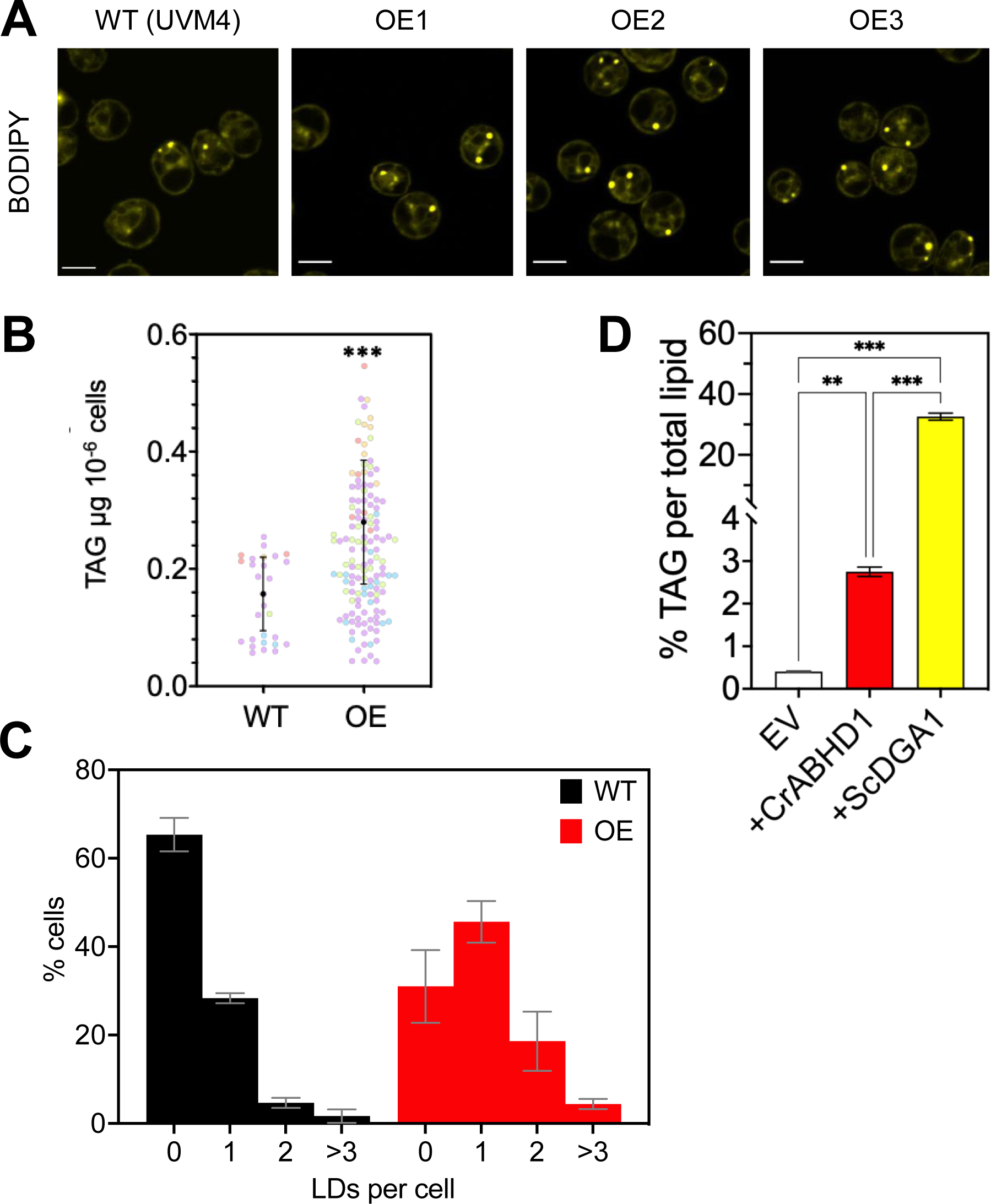
*ABHD1* over-expression boosts LD formation and TAG content. **A.** LD imaging in cells over-expressing *ABHD1* (Three independent lines are shown, bar = 5 µm). **B.** TAG amount in WT versus *ABHD1* overexpressing lines. Four independent experiments (color-coded differently) for six independent lines (three technical replicates for each line) are shown. Error bars show mean and standard deviation, Student’s t test: *** *p* <0.001. **C.** The distribution of LD numbers per cell analyzed (>200 cells examined per group). Histograms represent mean and standard deviation of three biological replicates for WT and five independent lines for OE (*ABHD1* overexpressor). **D.** TAG quantification in the H1246 yeast strain expressing the Chlamydomonas *ABHD1* gene. Yeast transformants expressing the respective gene were harvested and analysed for total TAG amount using GC-MS. Data are means of three biological replicates with standard deviation bars shown. ANOVA test: ** *p* < 0.01, *** < 0.001.

### Absence of ABHD1 affects the composition of the LD coat

To further investigate the function of ABHD1 in LD biogenesis, we isolated two insertional mutants from the *Chlamydomonas* mutant library (CLIP) (39). The two independent lines *abhd1-1* and *abhd1-2* harbor an insertion at the 3^rd^ and 10^th^ exon, respectively (**Fig. 3*A***). Using gene specific primers, RT-PCR analyses showed that there was no *ABHD1* expression in *abhd1-1* and *abhd1-2* mutant alleles (**Fig. 3*B***). Immunoblot analyses of total proteins extracted from isolated LDs from respective WT and the two mutant strains showed that no ABHD1 protein could be detected in either mutant (**Fig. 3*C***). Interestingly, the immunoblot from WT cells indicated that ABHD1 could be present as a dimer. Taken together, both gene expression and immunoblot analyses firmly confirmed both alleles as true knockout mutants. We then compared TAG amount in the two mutants (*abhd1-1* and *abhd1-2*) to their parental strain (i.e., CC4533) during N deprivation. No difference in TAG was observed under this condition (***SI Appendix*, Fig. S6*A***). Whole-cell lipidomics showed no significant change in membrane lipids either (***SI Appendix*, Fig. S6*B***). LD number and size are also similar between CC4533 and the two KO mutants (***SI Appendix*, Fig. S6*C*, *D***).

**Figure 3.**
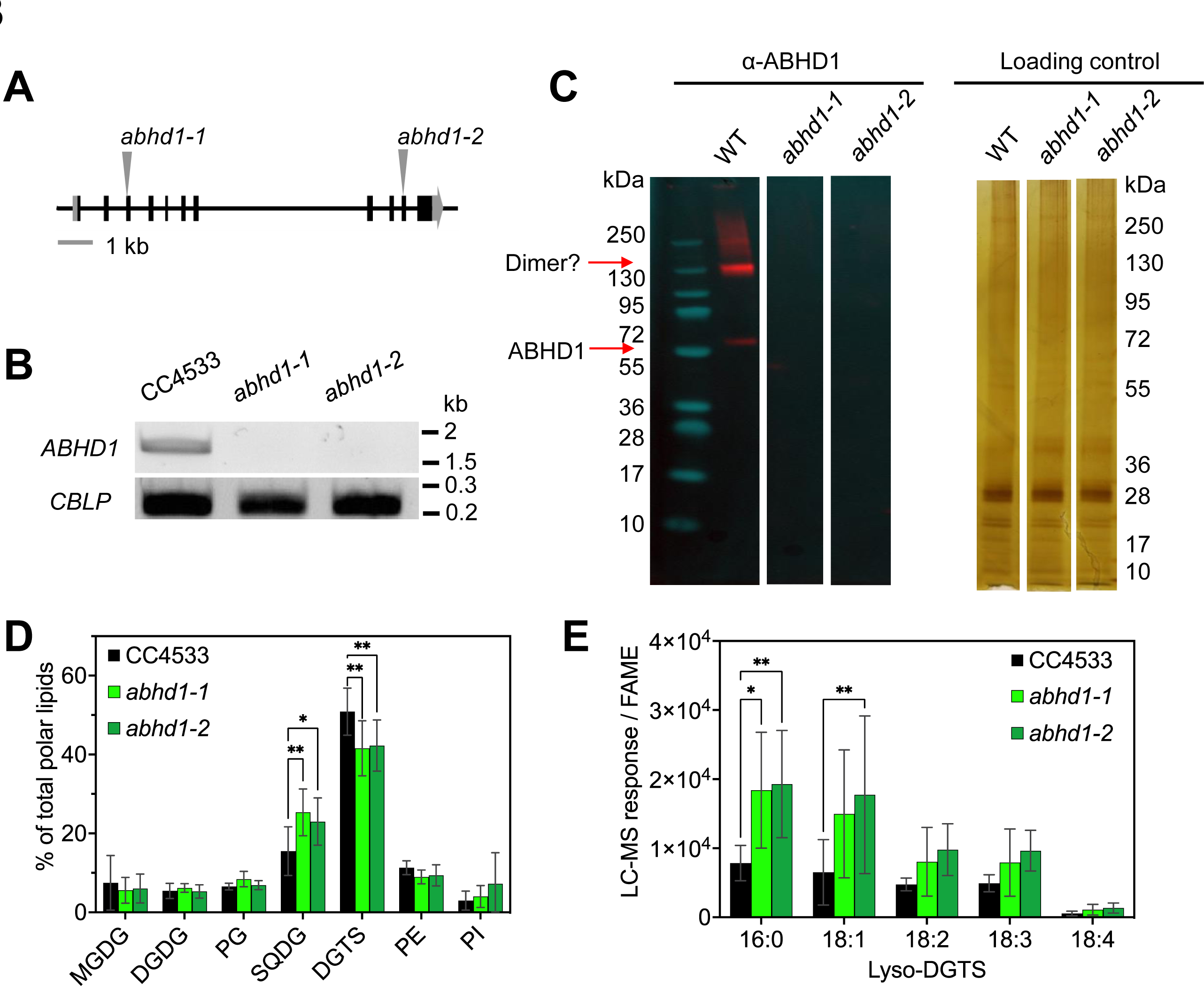
LD membrane coat analysis for WT and the two *abhd1-1* and *abhd1-2* mutants. **A.** The site of the *AphVIII* cassette insertion in the *ABHD1* gene for *abhd1-1* and *abhd1-2 mutants*. **B.** RT-PCR analysis of *ABHD1* transcript. **C.** Immunoblot analysis of ABHD1 protein. **D.** LD membrane coat composition. **E.** Changes in lyso-DGTS species in the mutants. LDs were extracted from cells N starved for 2 d. Proteins were extracted then loaded into each lane, based on a fixed number of fatty acid methyl esters (FAME) (i.e., ca. 55 μg FAME equivalent of LDs loaded). Silver-stained SDS-PAGE of LD-isolated proteins acts as a loading control. Levels of lyso-DGTS species detected in the polar lipid fraction of isolated lipid droplets. UPLC-MS/MS signal was normalized by FAME quantities in the polar lipid fraction. Data are means of five biological replicates from three independent LD isolations. Bars indicate standard deviation. P values were determined by a two-way ANOVA with Bonferroni correction: * *p* < 0.05; ** *p* < 0.01. Abbreviations: CBLP: *Chlamydomonas* beta subunit-like polypeptide; FAME, fatty acid methyl ester; MGDG, monogalactosyldiacylglycerol; DGDG, digalactosyldiacylglycerol; PG, phosphatidylglycerol; SQDG, sulfoquinovosyldiacylglycerol; DGTS, diacylglyceryl-*N, N, N*- trimethylhomoserine; PE, phosphatidylethanolamine; PI, phosphatidylinositol.

To investigate further the role that ABHD1 may have in LD formation in vivo, we performed comparative lipidomics and proteomics of isolated LDs from WT and the two knockout mutants. DGTS was identified as a major component of LD lipid-coat, followed by SQDG and PE. The mutants’ LD coat showed a remodeled lipid composition (**Fig. 3*D***), with a lower proportion of the major lipid class (DGTS) but a higher proportion of SQDG, a class of negatively charged lipids. Among the lipid species with the most significant differences were some lyso-DGTS species, in particular lyso-DGTS 16:0, which were present in significantly higher amount in the mutants’ LD coat compared to WT (**Fig. 3*E***).

To determine whether the mutants affected the flux of nascent fatty acids through membrane lipids into TAG during N-starvation, we performed a [^14^C]acetate pulse-chase analysis (***SI Appendix*, Fig. S7**). There were limited differences in the precursor-product relationships of membrane lipids and TAG between WT and mutant lines, indicating the action of ABHD1 does not affect the major flux of acyl chains into TAG during N starvation.

We explored potential proteomics changes in the isolated LDs (***SI Appendix*, Fig. S8**). Among the 598 proteins repeatedly detected by mass spectrometry (MS)-based proteomics, only four showed a significant change to the relative amount between the WT and the two mutants (**Dataset S1**). These four proteins were reduced in the mutant LDs: ABHD1 (as expected), the long-chain acyl-CoA synthetase (LCS2), the photosystem I reaction center subunit H (PSAH), and a putative glycosyl hydrolase (GHL1). The reduction in LCS2 protein amount in the two knockout mutants supports the idea that in the mutant there is indeed a reduced need for acyl-activation, corroborating the enzymology data below showing that ABHD1 is a lipase (next section). GHL1 has been reported to potentially function as a galactolipid galactosyl hydrolase and providing DAG for TAG synthesis (46), but it remains to be tested whether GHL1 participates in glyceryl-*N,N,N*-trimethylhomoserine (GTS) hydrolysis allowing therefore glycerol recycling downstream of ABHD1. Taken together, these data suggest that ABHD1 may act in the modification of the lipid membrane of the LDs as its absence results in changes in both LD’s membrane lipid and protein coat.

### ABHD1 has a lyso-DGTS lipase activity

To investigate the potential enzymatic activity of ABHD1, we expressed in *E. coli* a truncated version of ABHD1 lacking the N terminal hydrophobic portion (with amino acids 1 to 46 removed) resulting in recombinant ABHD1 (rABHD1). Purification of the soluble native rABHD1 was only partial but, in presence of urea, a highly pure rABHD1 protein was obtained from inclusion bodies and refolded (**Fig. 4*A**, Table S3***). To identify its potential substrate(s), the refolded purified rABHD1 was first incubated with total lipid extracts of *Chlamydomonas abhd1- 1* mutant. Interestingly, we observed a reduction in lyso-DGTS species, but no change in the other major lipid molecular species (**Fig. 4*B***). We could also detect changes in some other very minor lysolipid species (such as lyso-MGDG, lyso-SQDG, lyso-PG, lyso-PE) but not lyso- DGDG (***SI Appendix*, Fig. S9**). These other lysolipids were not further tested because they are > 100 times less abundant than lyso-DGTS. ABHD1 activity was then tested using pure lyso- DGTS as substrate prepared from the partial spontaneous hydrolysis of commercial DGTS. Results showed that ABHD1 indeed acted as a lipase on pure lyso-DGTS as the two expected co-products of the reaction (**Fig. 4*C***), palmitic acid and GTS, could be identified by MS (***SI Appendix*, Fig. S10**).

**Figure 4.**
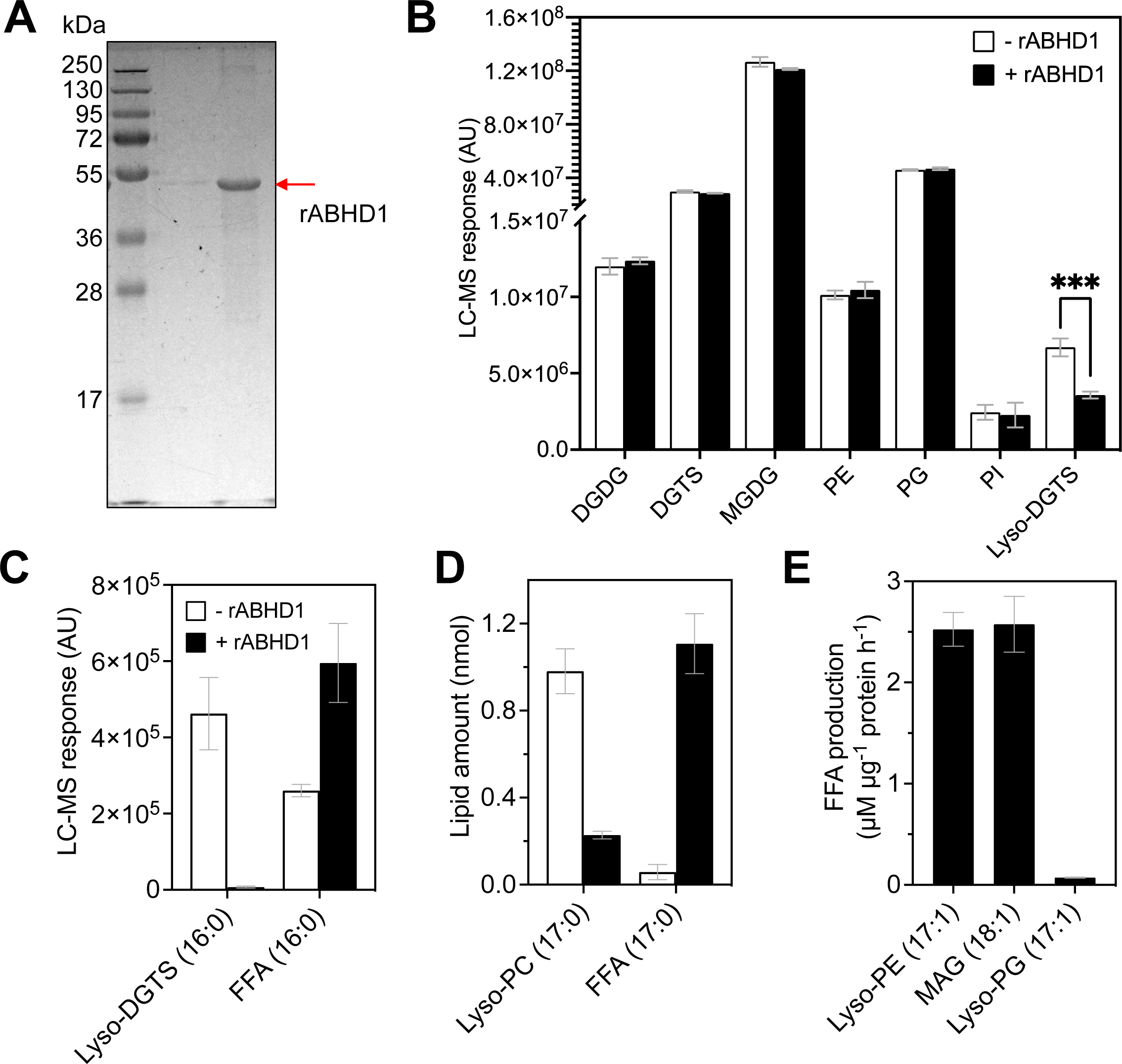
Enzyme activity analysis of the refolded rABHD1 protein. **A.** SDS-PAGE of rABHD1 purified from *E. coli* inclusions bodies and refolded. **B.** Activity test of rABDH1 on *abhd1-1* total lipid extracts identified lyso-DGTS as potential ABHD1 substrate. LC-MS/MS detected over 10000 m/z peaks for both the reaction with rABHD1 or without. **C.** The lipase activity of rABDH1 toward purified lyso-DGTS. **D.** The lipase activity of rABDH1 toward commercial lyso-PC. **E.** Activity assay on lyso-PE, lyso-PG, MAG, PC and DGTS. Data are the mean of three independent experiments and error bars refer to standard deviation. Student’s t-test: *** *p* < 0.001. Free fatty acid formation was quantified using 19:0 fatty acid as an internal standard. Reaction conditions for all ativity tests: Teorell Stenhagen universal buffer pH 7.5, with 100 mM NaCl and Triton-X100 at 1 CMC. During the enzymatic reaction (incubation of 2 h), during the preparation of lysolipid substrate or during lipid extraction, there is always some autolysis of the substrate, which is responsible for the presence of a certain amount of free fatty acids in the negative control. No activity could be detected on PC and DGTS for rABHD1.

Similar activity could be obtained using lyso-phosphatidylcholine (lyso-PC), a commercially available lipid structurally similar to lyso-DGTS (**Fig. 4*D***). Due to the difficulty in obtaining purified lyso-DGTS, further characterization of the putative enzymatic activity of rABHD1 was carried out on lyso-PC. Activity assays conducted under various conditions of buffer, pH, detergent or cations **(*SI Appendix*, Figure S11)** allowed to select a reaction medium for further studies. Release of free fatty acid from lyso-PC was found to increase with time and with rABHD1 amount, which showed that it corresponded to a genuine enzymatic activity (***SI Appendix*, Fig. S12**).

ABHD1 did not show activity toward lipids with two acyl-groups (**Fig. 4*B***), thus here we tested the activity of rABHD1 toward other lipids containing one fatty acid, e.g. lyso- phosphatidylethanolamine (lyso-PE), lyso-phosphatidylglycerol (lyso-PG), and monoacylglycerol (MAG). Results showed that rABHD1 protein released fatty acids from lyso- PE, MAG and lyso-PG but showed relatively low activity toward the latter (**Fig. 4*E***). This was observed on purified commercial substrates as well as when incubated with total lipid extracts (***SI Appendix*, Fig. S12**). These results thus indicated that rABHD1 has some preference for lysolipids over diacyl-lipid analogs (no activity toward lipids with two acyl-groups as shown in **Fig. 4*B***).

### ABHD1 is strongly inactivated by mutating the putative proton acceptor His356 to Ala

ABHD1 contains a putative lipase catalytic triad composed of a nucleophile (Asp179), an acid (Asp328) and a base (His356) residue (***SI Appendix*, Fig. S1*C***). The structural prediction of ABHD1 using alpha-fold (44) located these 3 residues of the putative catalytic triad close to each other on loop regions, thereby consolidating these features (***SI Appendix*, Fig. S13*A*).** To further demonstrate enzymatic activity of ABHD1, the histidine (H) at position 356 was mutated to an alanine (A), and the mutated protein was produced and purified from *E. coli* cultures. Activity was assayed using the commercial lyso-PC as a substrate, as ABHD1 was shown to possess a similar activity toward lyso-PC as lyso-DGTS. It was found that the H356A mutant had about 90% reduction in lyso-lipase activity compared to ABDH1 (***SI Appendix*, Fig. S13*B***).

### ABHD1 favours LD budding in vitro

We further investigated the effect of ABHD1 association and lysolipid hydrolysis on the geometry of LDs. To do so, we employed the droplet-embedded vesicle (DEV) system (47, 48). Model LDs are incorporated into pre-formed giant unilamellar vesicles (GUVs) producing DEVs (**Fig. 5*A***). We then deposited DEVs on a glass coverslip which caused them to rupture (**Fig. 5*B***), resulting in a 2D bilayer with the droplets incorporated (49). Resulting LDs were immobilized and their geometry was followed according to time. The phospholipid composition was as follows: dioleoyl phosphatidylcholine (DOPC)/Lyso-PC/Rhodamine-PE (79.5/20/0.5% w/w), and TAG to form neutral lipid droplets. Fluorescent lipid Rhodamine-PE acted as a membrane reporter and nitrobenzoxadiazole (NBD)-labelled TAG allowed us to visualize the lipid droplet. The model LDs’ initial shape is a spherical cap and addition of the purified rABHD1 resulted in the transition of the model LD from a flattened to a budded shape. This transition is shown by the fact that in the lipid bilayer the projected droplet surface area decreased with time by 40% as compared to the control, i.e., without any protein added (**Fig. 5*C***, **5*D***). Moreover, we obtained similar results with pure DOPC DEVs, i.e., in the absence of lyso-PC, the major substrates of ABHD1 (**Fig. 5*E*, 5*F***). Furthermore, when a non-refolded urea-purified form of rABHD1, which was enzymatically completely inactive (see Material and Methods), was tested in the DEV system, budding was still observed despite with slightly weaker capacity in comparison to the wild-type version of the protein (***SI Appendix*, Fig. S13*C***). Taken together, these results show that, independently of its lyso-DGTS lipase function, the binding of ABHD1 to the model LDs’ surface participates to droplet budding (47).

**Figure 5.**
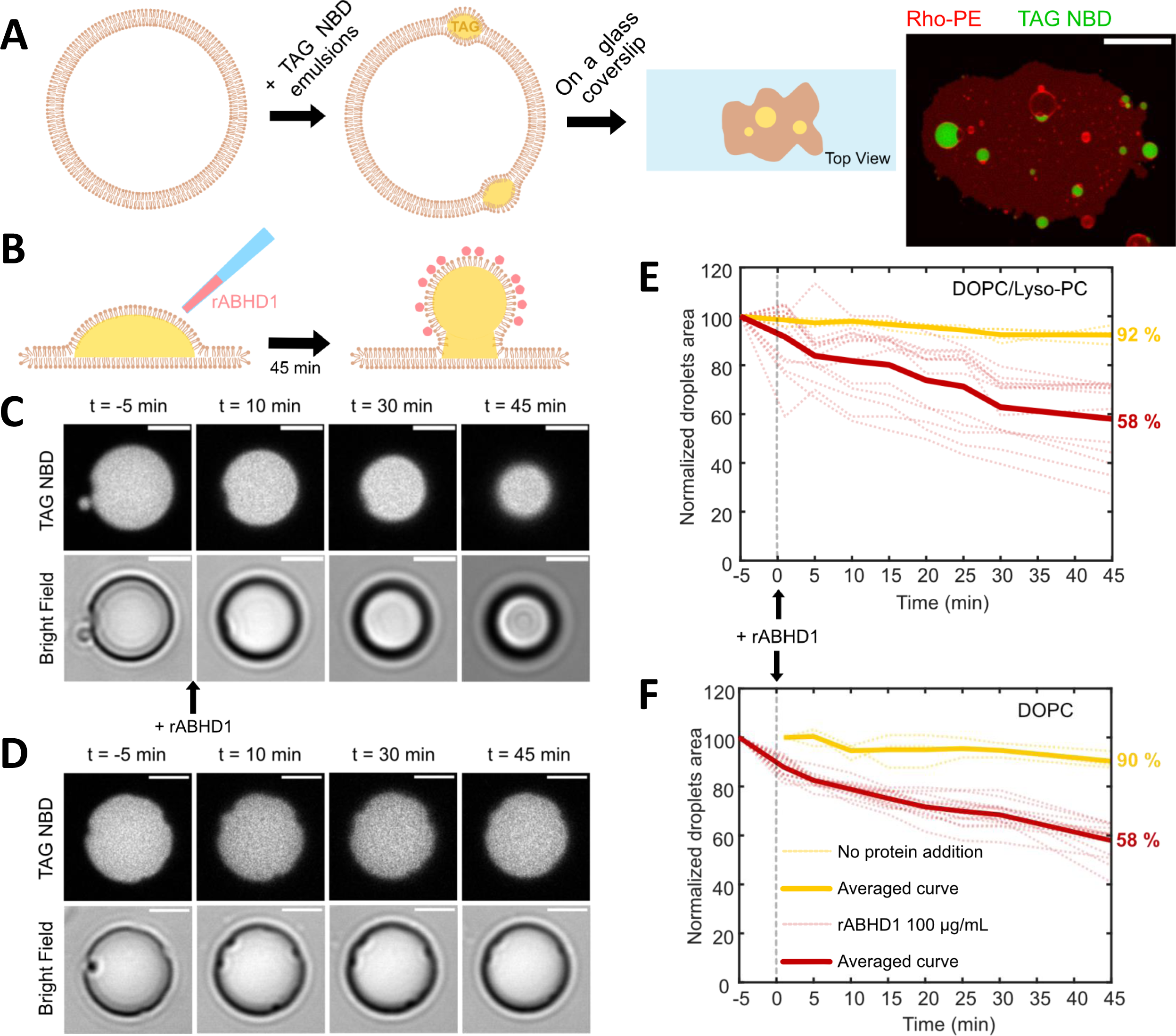
Addition of the rABHD1 protein to GUV boosts curvature formation. **A.** Schematic view of the experimental protocol. GUVs are put in contact with a thin emulsion of NBD-labelled triacylglycerol (TAG) that forms droplets at the lipid bilayer. When placed on a glass coverslip, membrane rupture results in a 2D lipid bilayer at the surface with droplets of TAG incorporated in. Typical image of a DEV after rupture is shown with lipids in red and TAG NBD in green. Scale bar is 10 µm. **B.** The droplet initially forms a spherical cap and is budding from the membrane 45 min after rABHD1 addition in the observation buffer (100 µg mL^-1^). **C.** Time lapse of a TAG droplet at the membrane plane (NBD TAG signal in upper panel and bright field in lower panel), showing the decrease of the surface after addition of rABHD1. Scale bars are 2 µm. **D.** Control time lapse of a TAG droplet at the membrane plane without addition of protein. Scale bars are 2 µm. **E, F.** Measurement of the droplet surface at the membrane plane according to time in two different lipid composition: (E) DOPC/Lyso-PC/RhoPE and (F) DOPC/RhoPE. The measured area of each droplet is normalized by the initial surface occupied at t = -5 min. Dashed lines represent the time traces of individual droplets and thick lines are the average trend. Yellow lines correspond to the control in the absence of ABHD1 [(E) N = 3 droplets analyzed, (F) N = 4], and red lines to addition of rABHD1 at 100 µg mL^-1^ [(E) N = 16, (F) N = 13].

## Discussion

Although the proteome of *Chlamydomonas* LDs was published more than 10 years ago (33, 34), only a few LD-associated proteins have been studied in any detail, and none have a demonstrated role in LD biogenesis. Here using a combination of biophysical, cell biological as well as genetic and lipidomic approaches, we show that the α/β hydrolase domain-containing protein ABHD1 acts as a lysolipid lipase and participates in LD biogenesis. Thus far, lysolipid acyltransferases have been studied in algae (50), but no lysolipid lipase has yet been identified. We demonstrate here that ABHD1 catalyzes the hydrolysis of an acyl group from a lyso-DGTS molecule, and it is therefore the first enzyme characterized to date acting as a lyso-DGTS lipase, and the first lysolipid lipase identified among algae. Our current working model is that ABHD1 promotes LD emergence likely through two actions at the LD surface: a lyso-DGTS lipase activity, and a distinct non-enzymatic property that promotes LD emergence, probably altering the metabolic fate of DGTS. We provide a model of molecular events involving ABHD1 that may lead to LD emergence (**Fig. 6**) and we then discuss the possible implications of our findings for DGTS and TAG metabolism.

**Figure 6.**
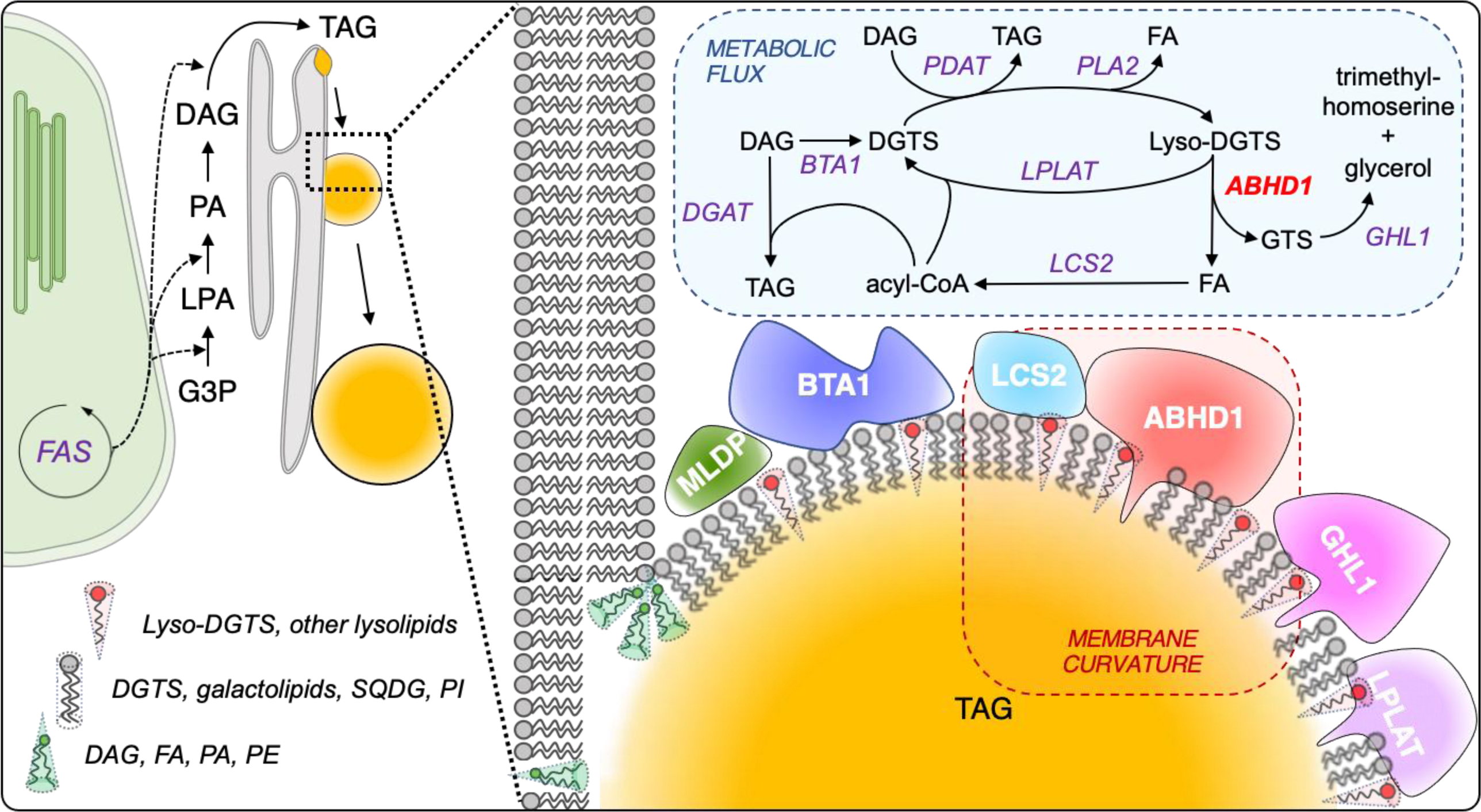
A model explaining the multifaceted actions of lyso-DGTS and ABHD1 protein in promoting LD budding, growth and TAG synthesis. ABHD1 catalyzes the hydrolysis of lyso-DGTS to produce a free fatty acid and a GTS moiety. ABHD1 is present at basal levels during optimal growth, and it increases in transcription and protein amount as N starvation is initiated, paralleling the increases in both LD number and size (64). During LD budding, the formation of lyso-DGTS and the synthesis of ABHD1 protein both promote positive membrane curvature therefore LD budding. Later on, ABHD1 activity may facilitate LD growth as less curvature is needed by degrading lyso-DGTS. The resulting FA can be activated to acyl-CoA by LCS2 and contribute to TAG synthesis. Additionally, by consuming lyso-DGTS, ABHD1 is pulling the flow of acyls from either a putative PDAT or a PLA2, whose activity drives either TAG or FA production, respectively. Indeed, in two knock-out mutants, LCS2 protein is present in reduced amount, likely because the flow of free FA is reduced without ABHD1.Taken together, the major role of ABHD1 lies in LD coat remodeling and maintain LD stability. Abbreviations: BTA1, betaine lipid synthase 1; DAG, diacylglycerol; DGAT, diacylglycerol acyltransferase; DGTS, diacylglyceryl-3-O-4’-(*N,N,N*-trimethyl)-homoserine; FAS, fatty acid synthase; G3P, glycerol-3-phosphate; GPAT, glycerol-3-phosphate acyltransferase; MLDP, major lipid droplet protein; LCS2, long chain acyl-CoA synthetase; Lyso-PA, lysophosphatidic acid; LPLAT, Lysophospholipid acyltransferase; PA, phosphatidic acid; PDAT, phospholipid:diacylglycerol acyltransferase; PLA2, phospholipase A2; TAG, triacylglycerol.

### ABHD1 is a lyso-DGTS lipase acting on LD surface

ABHD1 belongs to the α/β-hydrolase fold domain-containing superfamily, one of the largest protein superfamilies, present in virtually all sequenced genomes (51). Despite their large number and wide occurrence, the physiological substrates and products for members of the ABHD family remain largely uncharacterized (51). Here through in vivo and in vitro experiments, we provide firm evidence that ABHD1 is located at the LD surface (**Fig. 1**) and is a novel type of lipase, namely a lyso-DGTS lipase (**Fig. 3**). ABHD1 is indeed shown to be active in vitro on lyso-glycerolipids, including lyso-DGTS, but not on diacyl-glycerolipids such as DGTS. Similar to some other members of its superfamily, ABHD1 contains the lipase catalytic triad, i.e., Asp179, Asp328 and His356 (***SI Appendix*, Fig. S13*A***). We show here in this study that H356 is important for its catalytic activity because when it is replaced by an Ala, 90% of the lyso- lipase activity is lost. Furthermore, the orientation of the protein on the LD surface could be favored by the C-terminal hydrophobic helix (***SI Appendix*, Fig. S1*B***), bringing the putative active site of the enzyme closer to the LD surface.

### Lysolipids and biophysical properties of ABHD1 contributes to LD budding

Independent to its enzymatic function, ABHD1 is also found to play a structural role in favoring LD budding and growth (based on both in vivo and in vitro experiments (**Fig. 2**, **Fig. 5, SI *Appendix* Fig. S13**). Lipidomics of isolated LDs from the two knock-out mutants show an increase in lyso-DGTS. Lysolipids contain a big polar head and only one acyl chain, and in general are present as hexagonal tubes I (HI) and form positively curved monolayers (52, 53). This property could be important for the budding process because positive curvature forms sharp angles. Indeed, higher amount of lyso-DGTS was present in LDs of the two mutants. We therefore propose that at the initiation of LD formation, DGTS is converted to a lyso-DGTS which is important to facilitate membrane budding. In addition, a new study has recently reported that lyso-DGTS enhances the activity of PON1 (Paraoxonase 1), by so doing preventing oxidation of low-density lipoproteins (LDL)(54). This study therefore points out two potential important roles for lyso-DGTS, i) boosting enzyme activity by perhaps changing LD biophysics; ii) possible medical applications.

Nevertheless, the accumulation of lysolipids is deleterious to the ER membrane integrity (55). Therefore, while the generation of lyso-DGTS from DGTS may facilitate LD nucleation and budding, it will be crucial to prevent the accumulation of lysolipids. This could be done by balancing the actions of DGTS-to-lyso-DGTS conversion and lyso-DGTS degradation (**Fig. 6**). ABHD1 may mediate the specific degradation of lyso-DGTS (and possibly other lysolipids of the LD) by releasing free fatty acids from lysolipids. These fatty acids are then possibly used to make TAGs. A question remains as to why this reaction occurs on LDs: very likely because the LD surface offers more accessibility to lipids, as it has more packing defects (8, 56), which might grant better access to lyso-DGTS and their hydrolysis/detoxification. Knowing that the ER and LDs are in contiguity, lysolipids can diffuse between these organelles and their degradation at LDs would behave as a thermodynamic pump, driving lyso-DGTS from the ER-to-LDs for their continuous degradation.

### DGTS metabolism and TAG synthesis at the LD surface

Chlamydomonas lacks PC and contains instead the betaine lipid DGTS as its major extra-chloroplast membrane lipid (57) and lipid of LD surface (33). Compared to the large variety of proteins known to be involved in acylglycerol lipid metabolism (58), only one protein (i.e., BTA1) of betaine lipid metabolism has so far been studied, first in *Rhodobacter sphaeroides* (59), then in *Chlamydomonas* (38), and recently in *Nannochloropsis oceanica* (60). BTA1 catalyzes the addition of S-adenosylmethionine to diacylglycerol (DAG) forming DGTS (38). Recently, it has been shown that DGTS remodeling contributes to TAG synthesis during ER stress and this process is regulated by the transcription factor bZIP1 in *Chlamydomonas* (23). Pulse-chase labeling experiments revealed that DGTS turnover could provide acyl groups for TAG biosynthesis, however at a much lower level than MGDG or DGDG, similar to other previous results in *Chlamydomonas* (61, 62). This explains the lack of flux of ^14^C-acyl groups from DGTS to TAG in the *abhd1* mutants, consistent with the finding that its role is not in the major TAG biosynthetic pathway, but rather in the DGTS remodeling at the LD surface. Interestingly, BTA1 together with ABHD1 is found to be among the top 10 most abundant proteins in LDs from WT cells (**Dataset S1**), indicating that the LD surface provides a platform for active DGTS synthesis and turnover that may be separated from roles of DGTS in the ER membrane. The role of ABHD1 in LD coat remodeling and TAG synthesis is further supported by the changes in protein compositions observed in the two knock-out mutants (**Fig. 6**).

Taken together, the synthesis and degradation of lyso-DGTS at the LD surface indicates ABHD1 plays a dual role to initiate LD formation and ensure LD growth, important in adapting membrane biophysical properties as well as in supplying acyl-chains for TAG synthesis. The fact that the knock-out mutants (*abhd1-1* or *abhd1-2*) did not show differences in their TAG content suggests that ABHD1 does not constitute a major flux in supplying acyl-chains for TAG synthesis, but rather it is important in membrane lipid coat remodeling and in preparation of biophysical membrane properties essential for LD emergence and assembly. Overexpression of the ABHD1 protein during N-replete optimal growth boosted LD formation and oil content in Chlamydomonas but also in yeast **(Fig. 2)**, suggesting that ABHD1 could be used to increase TAG content of microalgae, and possibly other organisms, without compromising/halting cell growth.

## Materials and methods

### Strains and culture conditions

All *Chlamydomonas* strains were maintained in agar plates containing Tris-Acetate-Phosphate (TAP) media (63) under constant light at 25°C. In the case of mutants, TAP agar plates were supplemented with 15 μg mL^-1^ hygromycin or 10 μg mL^-1^ paromomycin as appropriate. Cell cultures were grown in TAP liquid media in conical glass flasks kept in incubators (Multitron, Infors HT) shaking at 120 rpm, 25°C, under continuous light (80-100 µmol photons m^−2^ s^−1^). Cell concentration, size and volume was quantified with a Multisizer 4 (Beckman Coulter). To start N deprivation, log-phase cultures were washed with fresh media without N (NH4Cl replaced by an equal molar amount of NaCl) at 450 *g* for 3 min.

All other methods are described in ***SI Materials and Methods*.**

## Acknowledgements

ITR is recipient of an international PhD studentship from CEA (program Irtelis). We thank Stéphanie Blangy and Pascaline Auroy in providing guidance on protein expression and molecular biology work respectively. We also thank Ousmane Dao in helping with culture preparations and Pascale David for verifying some constructions. We thank Audam Chhun for helping with graphic design. We also acknowledge the European Union Regional Developing Fund (ERDF), the Région Provence Alpes Côte d’Azur, the French Ministry of Research and the CEA for funding the HelioBiotec platform. This work is also partly supported by DRF Impulsion LD power. We thank the ZoOM Microscopy facility (BIAM, CEA Cadarache) and thank the Chlamydomonas Resource Center at the University of Minnesota for providing the indexed *C. reinhardtii* insertional mutants. A.R.T. is supported by ANR-21-CE11-0032-02- LIPRODYN. M.G.G, H.K.K, P.D.B. are supported by National Science Foundation PGRP-IOS- 1829365. The proteomic experiments were partially supported by Agence Nationale de la Recherche under projects ProFI (Proteomics French Infrastructure, ANR-10-INBS-08) and GRAL, a program from the Chemistry Biology Health (CBH) Graduate School of University Grenoble Alpes (ANR-17-EURE-0003). We thank the financial support of a collaborative project between Chinese Academy of Science and CEA.

## Supporting Information

**Figure S1.**
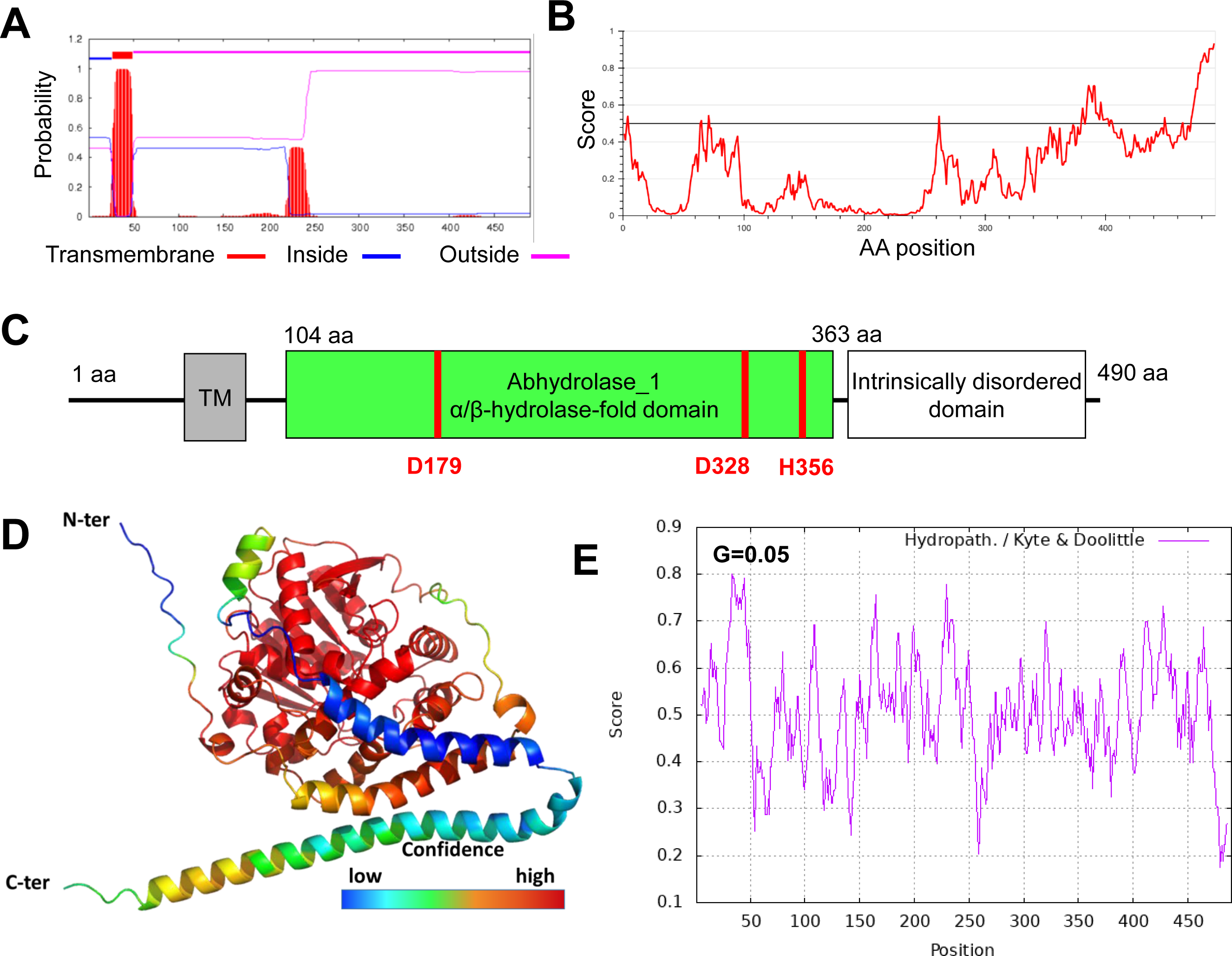

**Figure S2.**
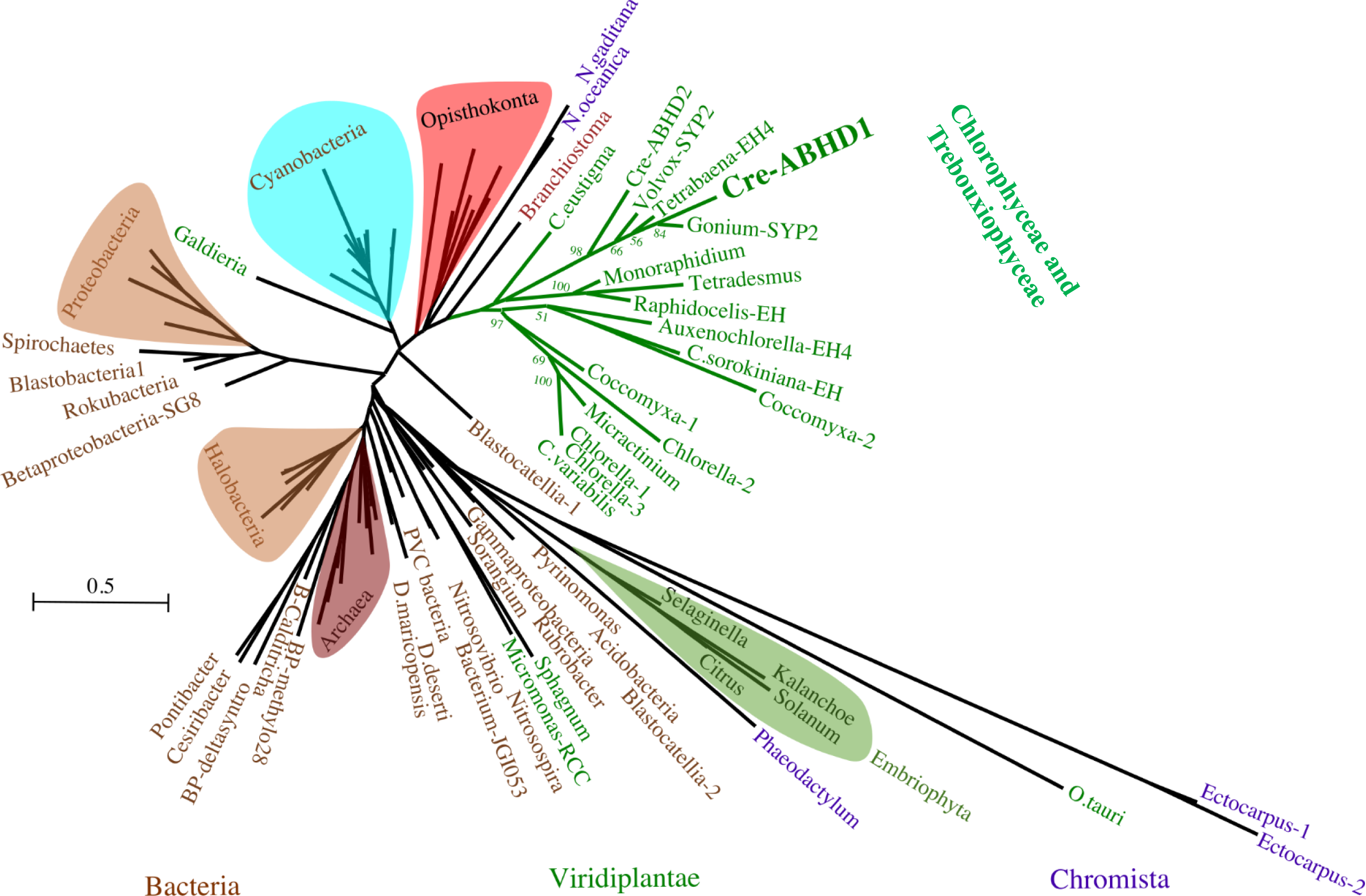

**Figure S3.**
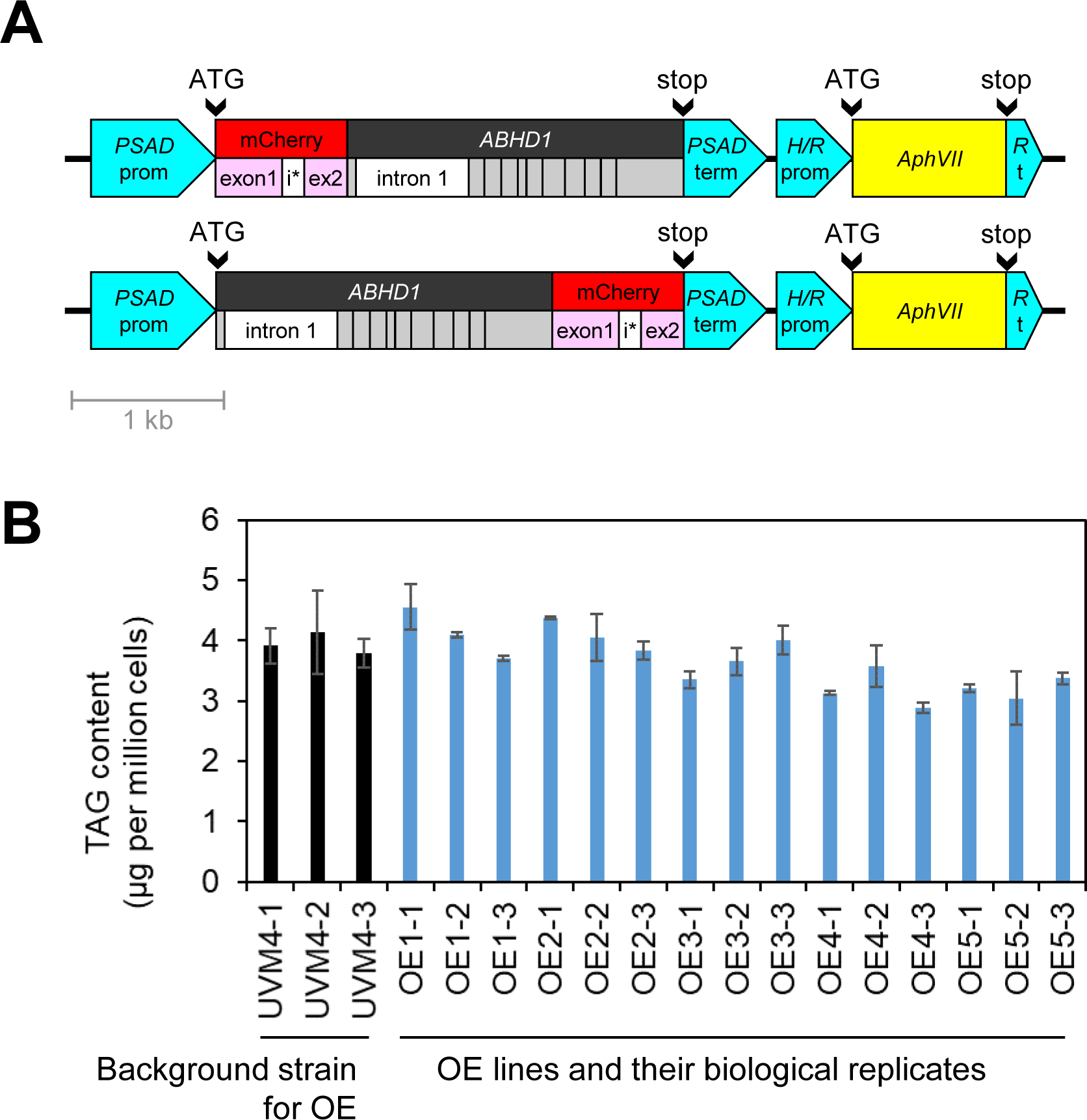

**Figure S4.**
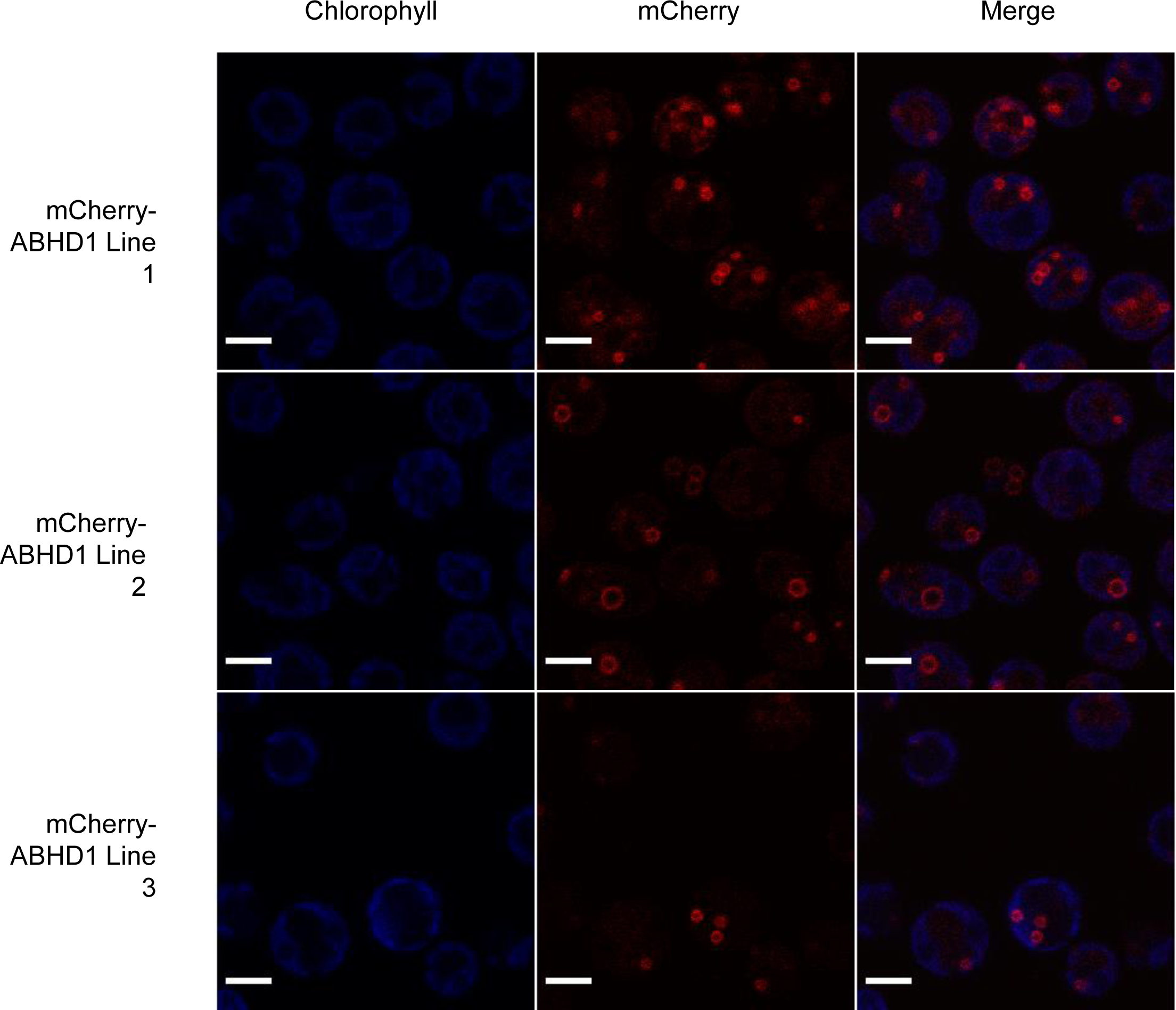

**Figure S5.**
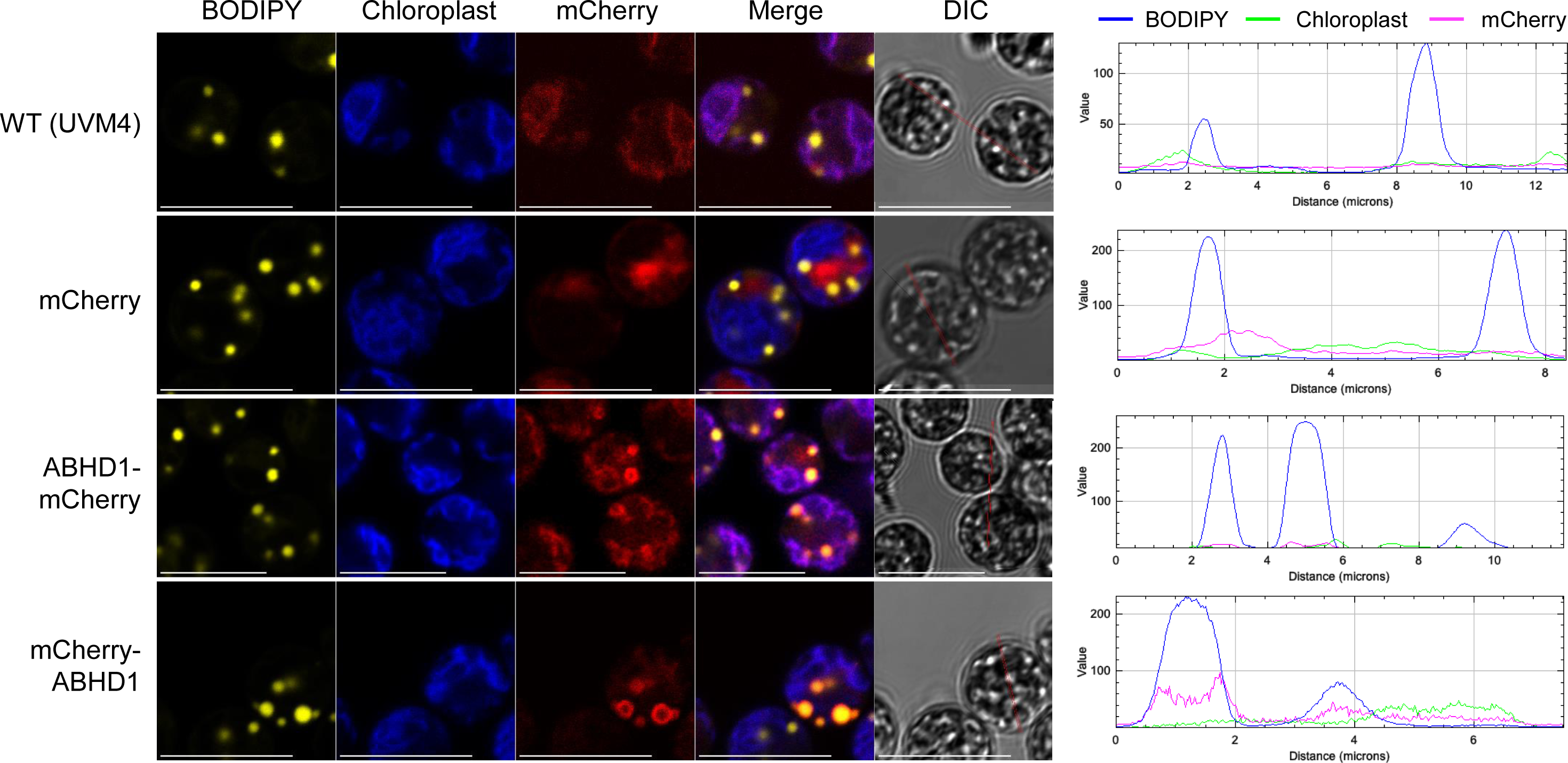

**Figure S6.**
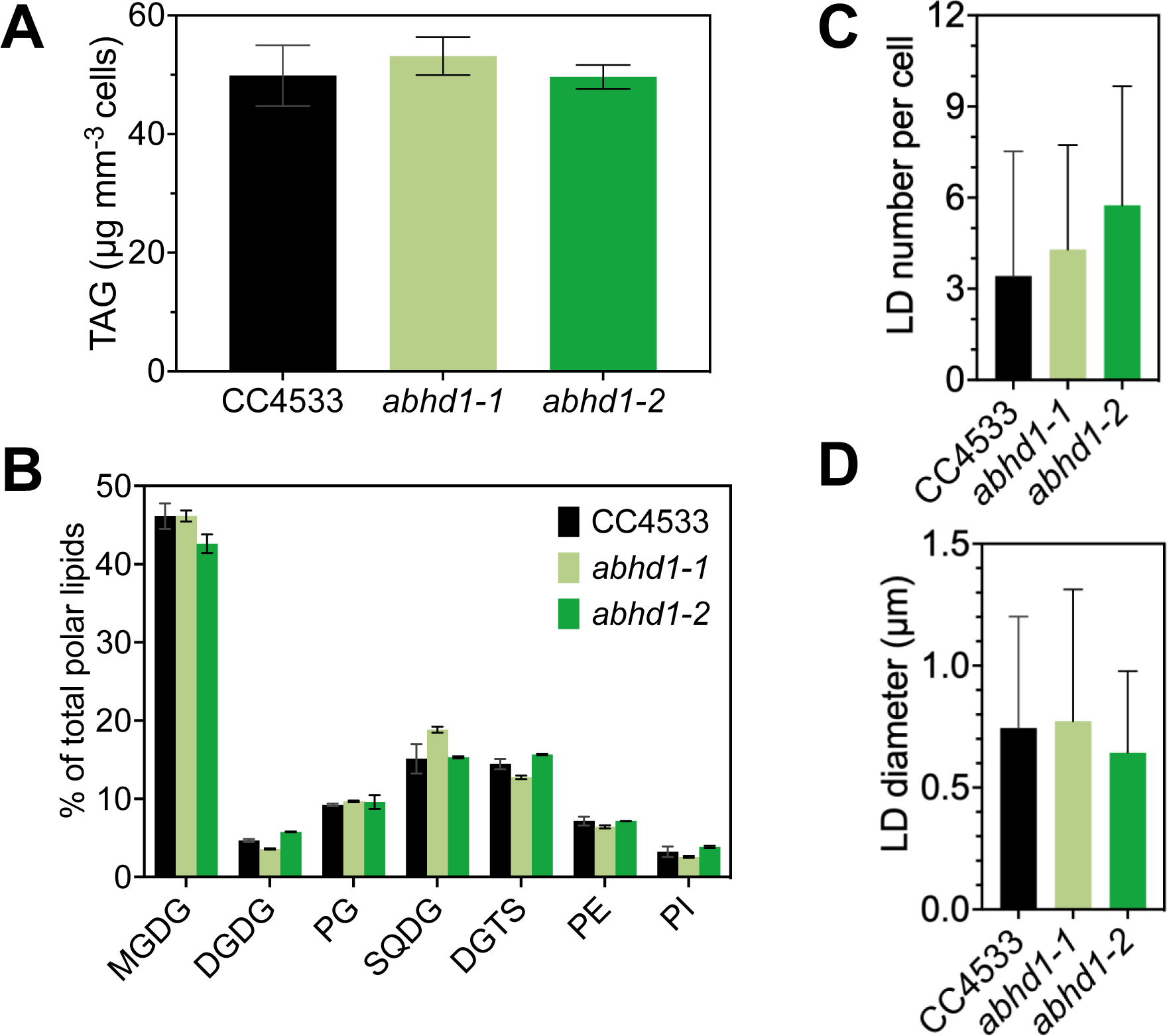

**Figure S7.**
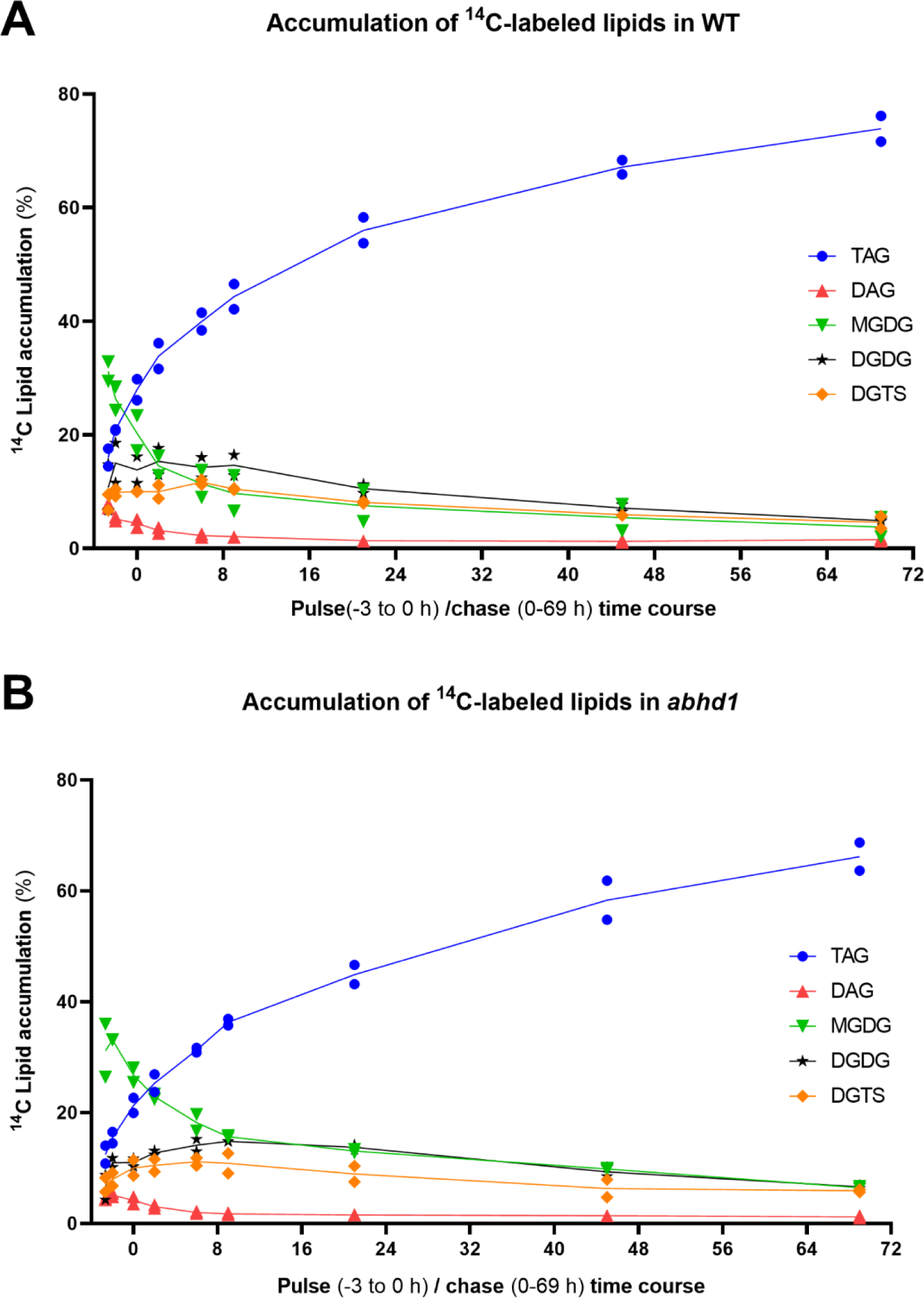

**Figure S8.**
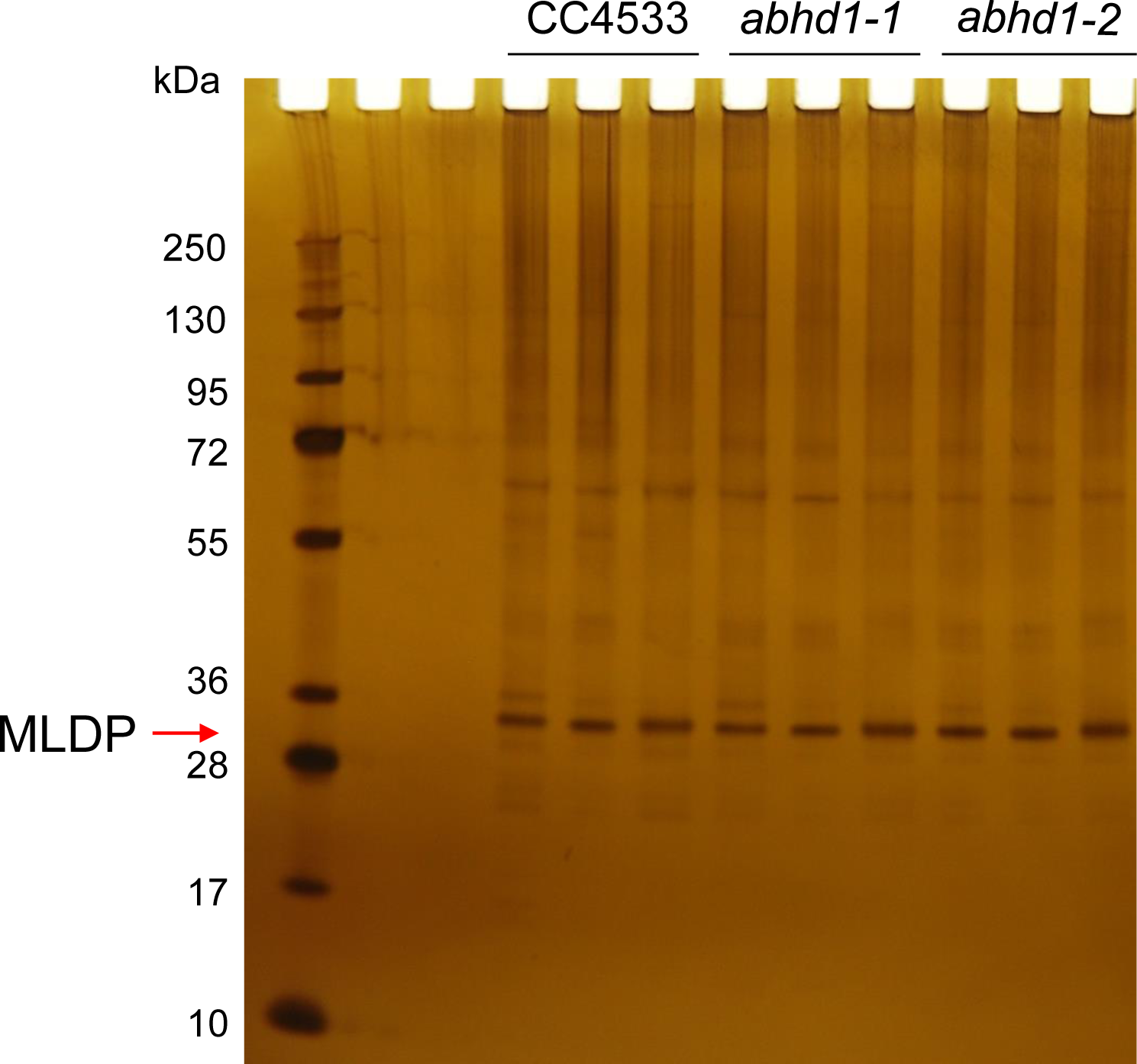

**Figure S9.**
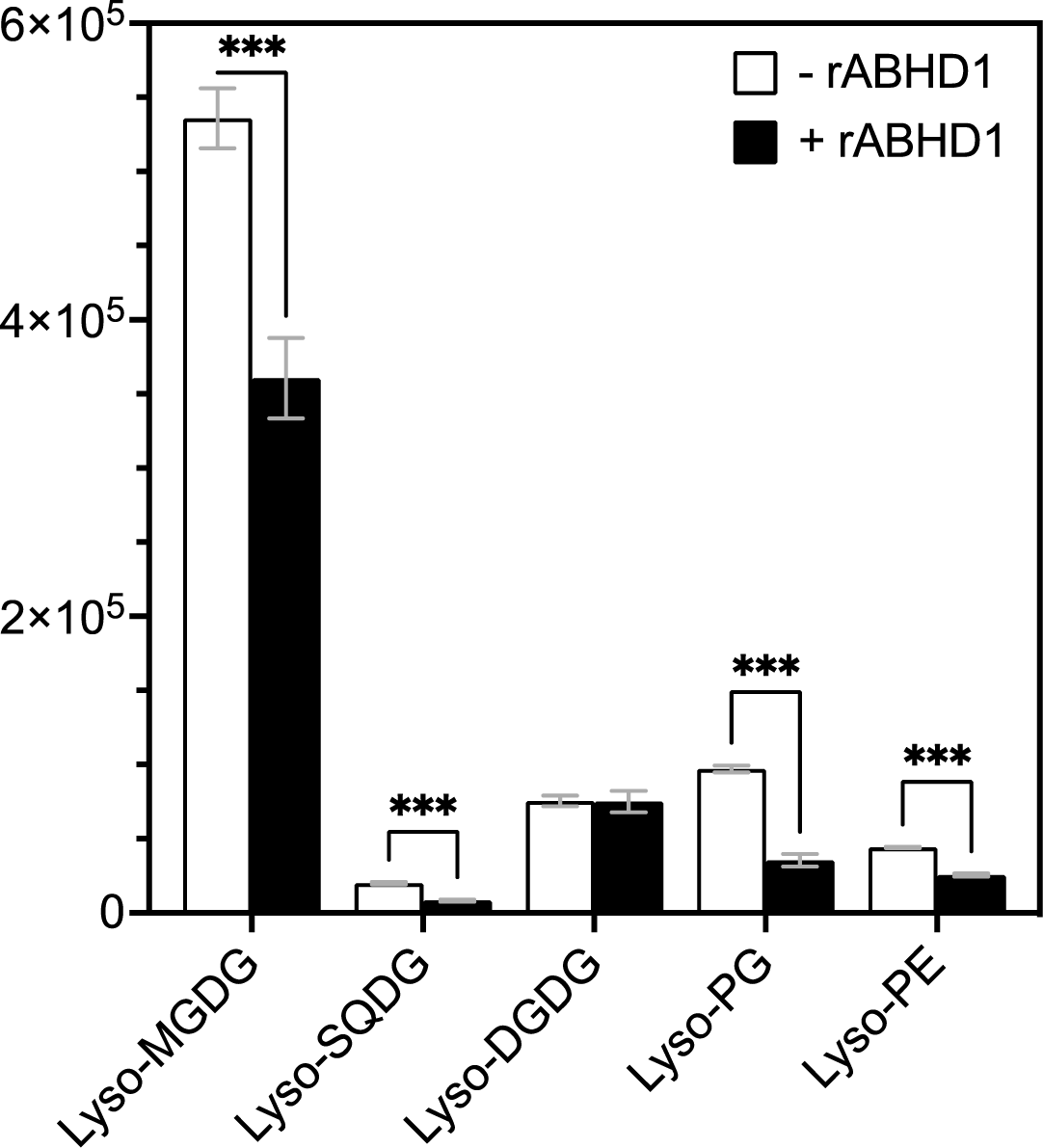

**Figure S10.**
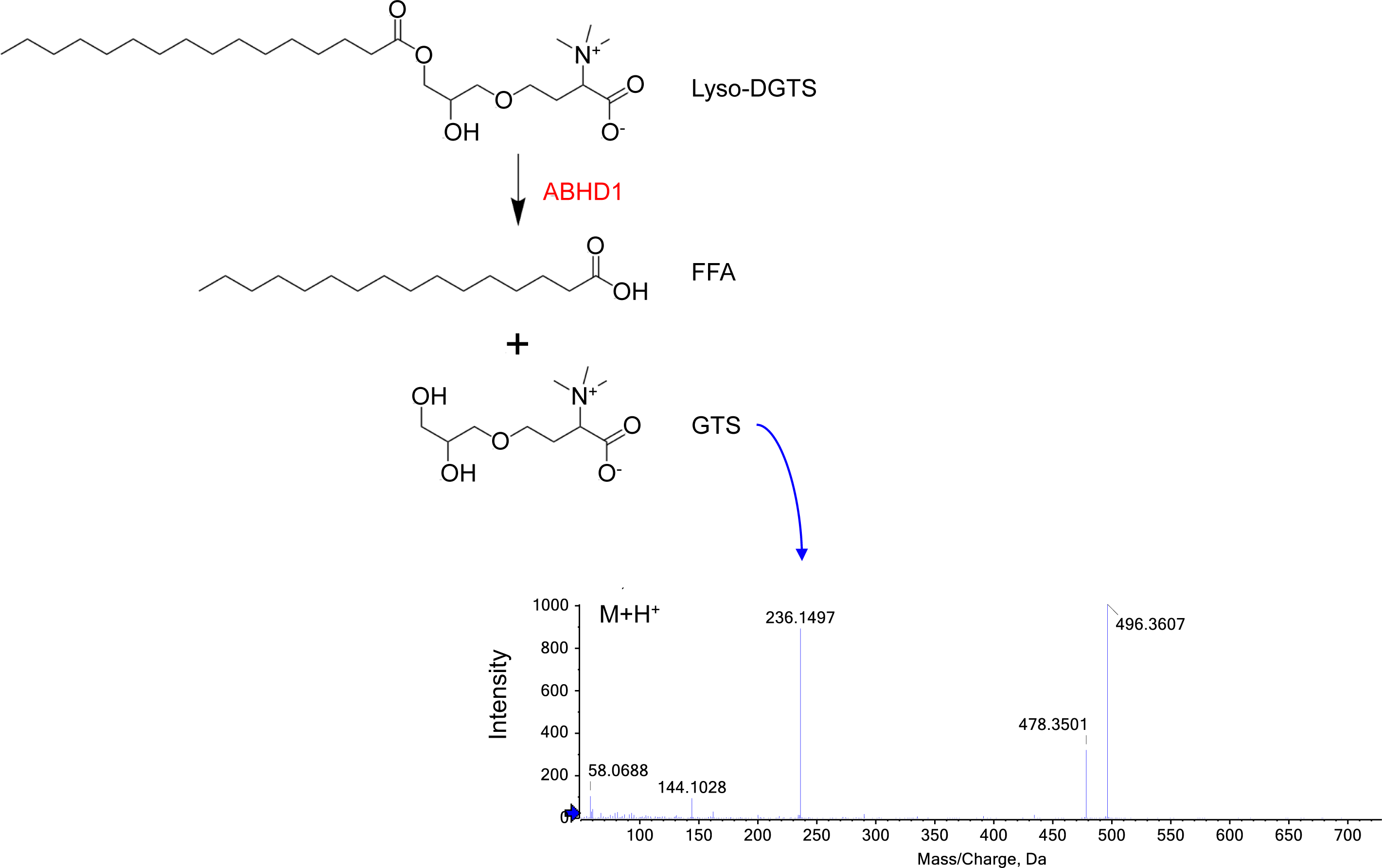

**Figure S11.**
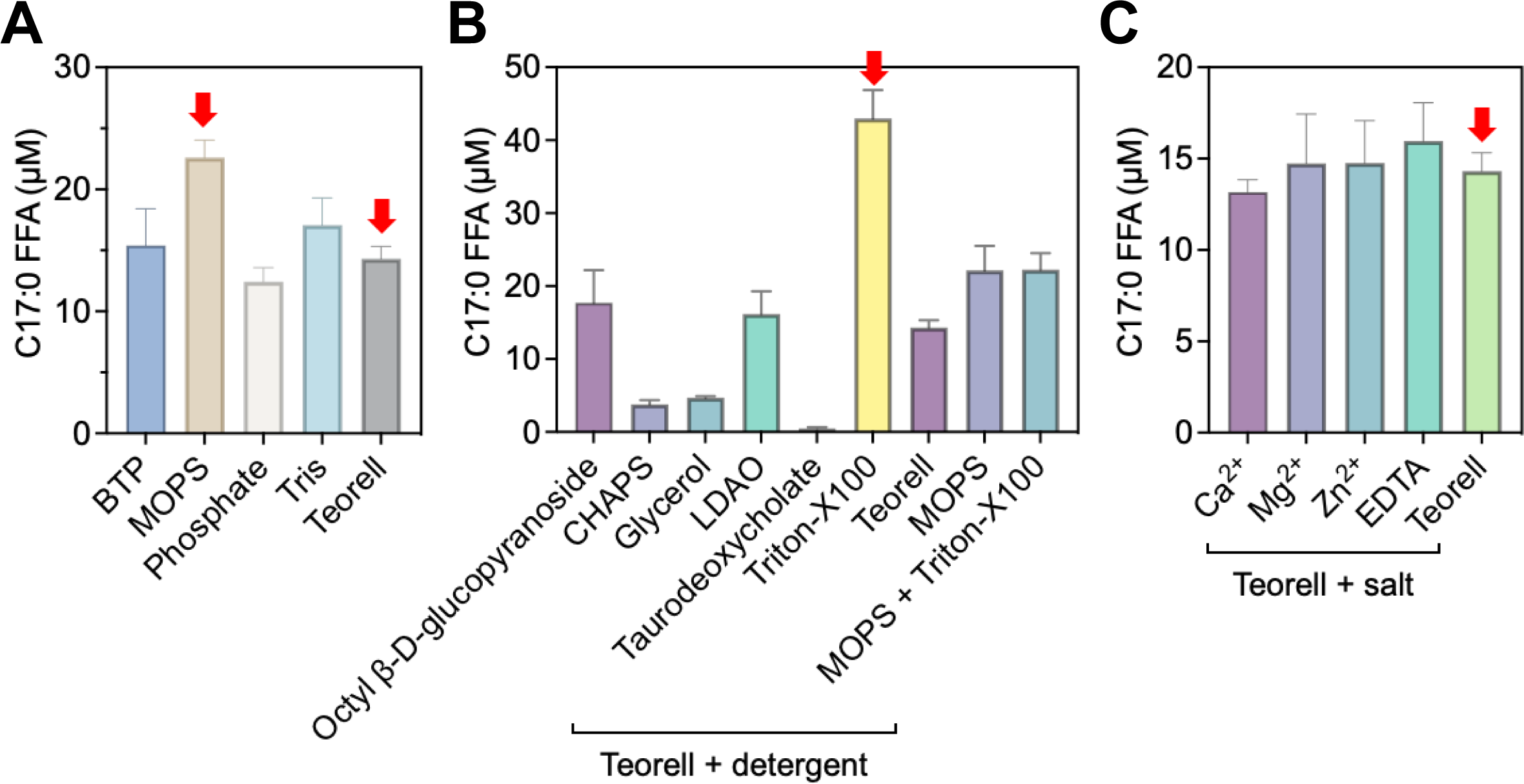

**Figure S12.**
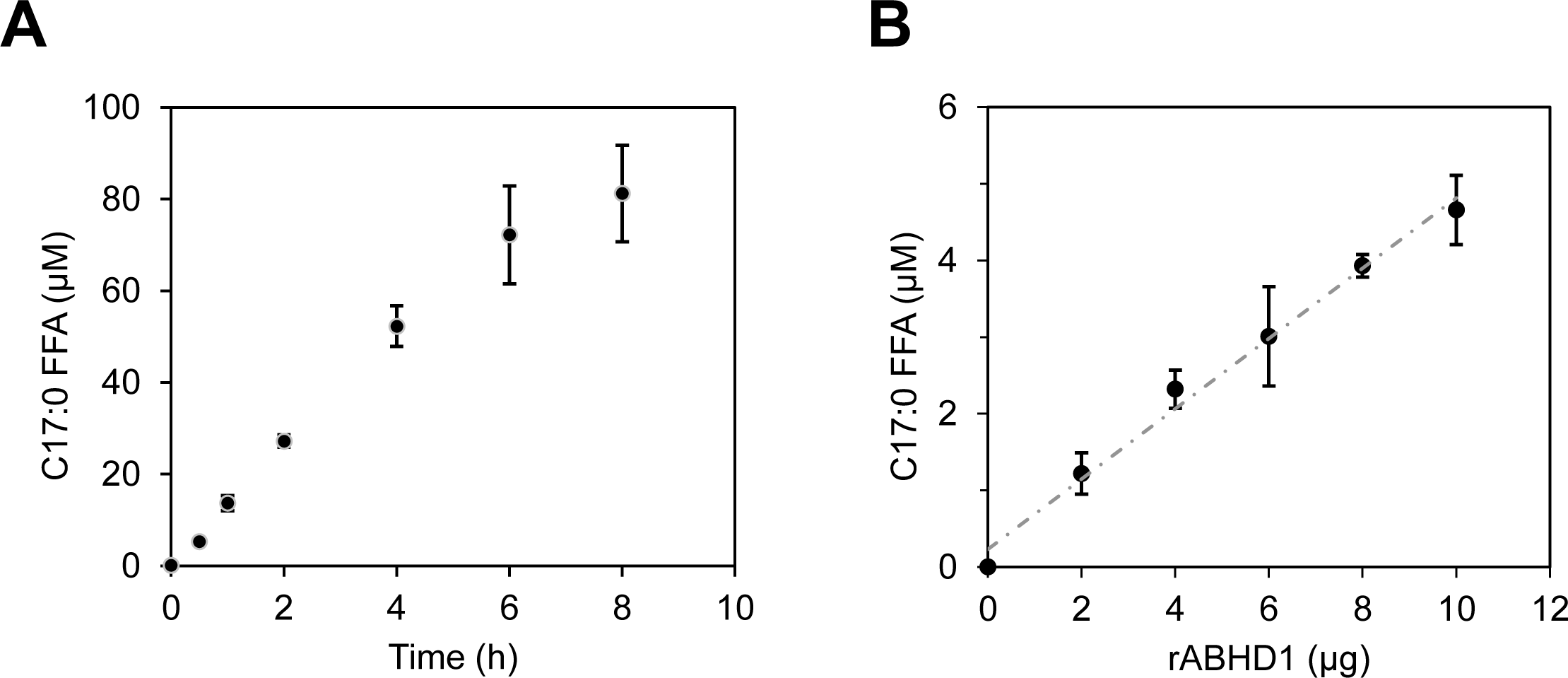

**Figure S13.**
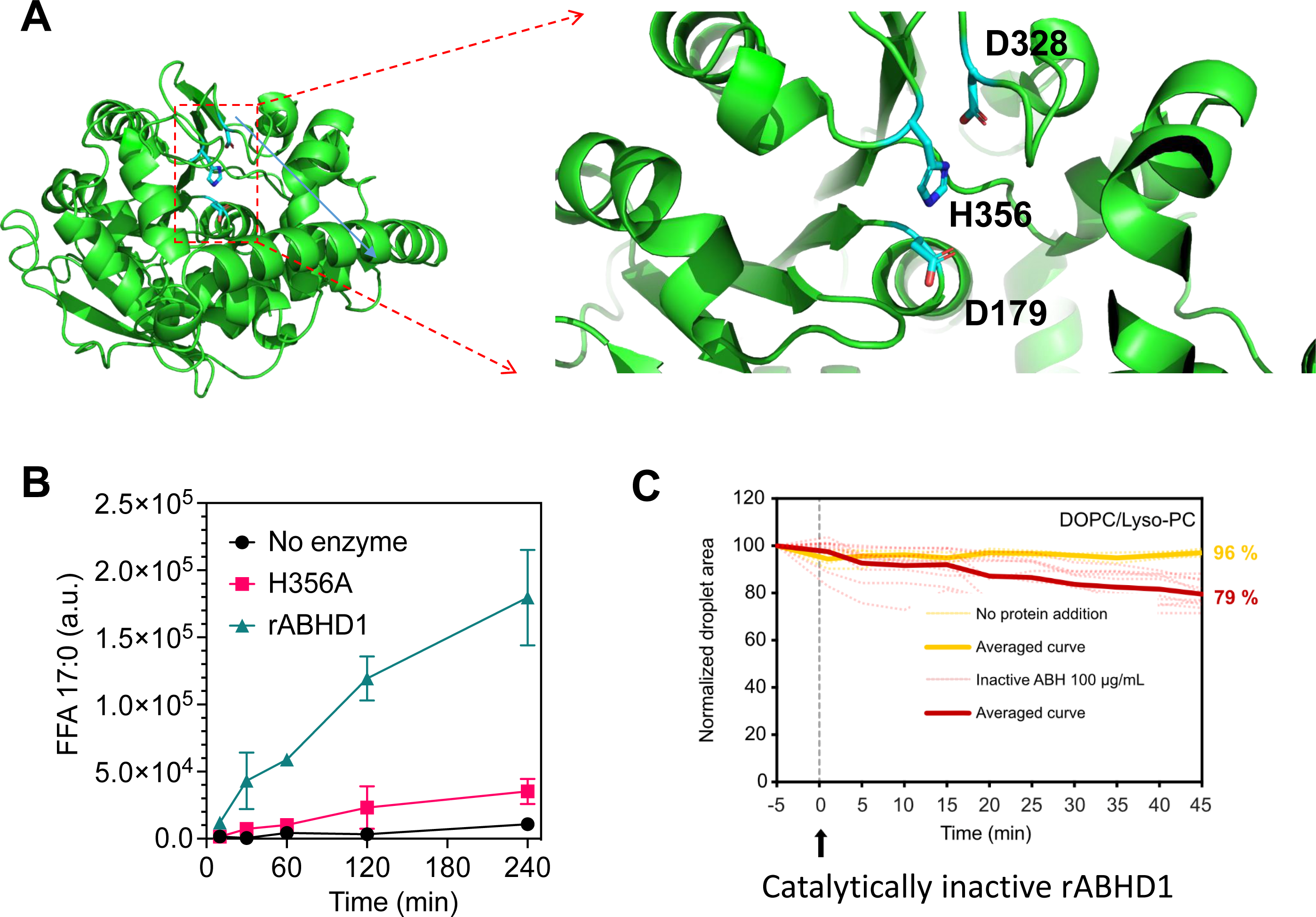

## Notes

### Competing Interest Statement

The authors have declared no competing interest.

