## Supplementary figures and images for "The α/β hydrolase domain-containing protein 1 (ABHD1) acts as a lysolipid lipase and is involved in lipid droplet formation"

### movie s1

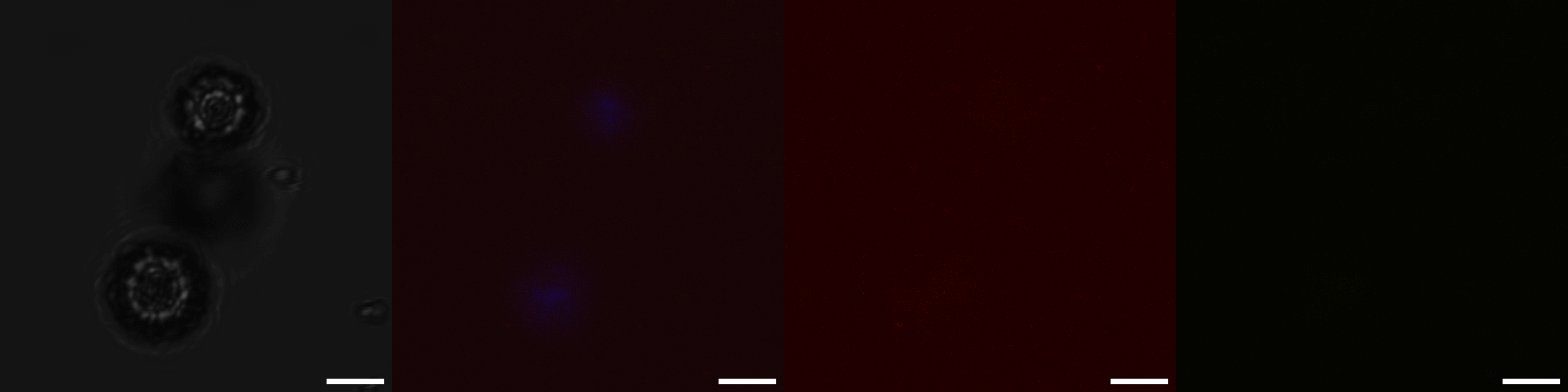
