## supplemental files for "The α/β hydrolase domain-containing protein 1 (ABHD1) acts as a lysolipid lipase and is involved in lipid droplet formation"

**This PDF file includes:**

SI Materials and methods
SI Figures S1-S13
SI Tables S1-S3
Movie S1
Dataset S1
SI References

### Materials and Methods

#### Bioinformatics

Sequences were first searched using BLAST tools. Hydropathy plot was generated using the Kyte-Doolittle algorithm (1). The value G in the graph is the grand average of hydropathy (GRAVY) value for each protein and was calculated using the GRAVY calculator (https://gravy.laborfrust.de/). IUPred3 was used to predict the disordered protein regions at https://iupred.elte.hu/ (2). TMHMM (TransMembrane prediction using Hidden Markov Models) was used to predict potential transmembrane domains (3). 3D structural features were predicted using α-fold algorithm (4) and then visualized using PyMOL.

#### Phylogenetic analysis

ABHD1 protein sequence was blasted on all plant and algal species from JGI Phytozome 12, fifteen algal species from Genome Portal, and other species from UniProtKB databases. A dataset of 285 sequences was curated to remove repeated sequences with low E-values and have a balanced representation of every clade. Finally, sequences of 92 proteins were aligned in MAFFT v.7 and visualized in Seaview program (***SI Appendix*, Table S1**). Phylogenetic trees were generated using Gblocks, protein regions highly conserved, by the maximum likelihood algorithm PhyML 3.1 (5) setting 100 as bootstrap. Seaview circular unrooted tree was edited.

#### Verification of *ABHD1* expression level in the *abhd1-1* and *abhd1-2* mutants

Two insertional mutants LMJ.RY0402.130997 (*abhd1-1*) and LMJ.RY0402.186121 (*abhd1-2*), and its genetic background CC4533 were purchased from the CLIP collection (https://www.chlamylibrary.org/) (6). To verify the level of expression of *ABHD1* gene, total RNA was extracted from mid-log phase cultures (at around $5\times10^{6}$ cells $mL^{-1}$). RNA extraction was performed as detailed in (7). Yield was checked by Nanodrop and 1 µL of total RNA was retro-transcribed with the cDNA synthesis kit SuperScript III First-Strand Synthesis SuperMix for qRT-PCR (Invitrogen). To amplify the *ABHD1* coding sequence (1473 bp, locus Cre12.g540550), KOD Xtreme Hot start polymerase (Novagen) was used with the primers ABH1_E1_F and ABH1_E11_R. The G protein beta subunit-like polypeptide (*CBLP*) gene (279 bp, Cre06.g278222) was used as a housekeeping gene (8), and amplified using primers CBLP_F and CBLP_R. These and all other primer sequences used in this study are given in ***SI Appendix*, Table S2**.

#### Plasmid construction for protein subcellular localization

It is well known that transgene expression can be enhanced by including the first endogenous intron (9). We therefore cloned a hybrid version of the gene for *ABHD1*. The first half of the cloned gene includes the genomic DNA covering 728 bp upstream region of the 5'UTR to the 2$^{nd}$ exon. The second half of the gene includes only the coding region from the 3$^{rd}$ exon to the 3'UTR. The total gene length is 3.4 kb including 700 bp up stream region (5'UTR), exon 1, intron 1, then followed by the rest of exons (exons 2-11, i.e. cDNA) and also 3'UTR (***SI Appendix*, Fig. S1**). Total genomic DNA was extracted from exponentially growing cells as described in (10). Primers ABH1-GP-F and ABH1-GE2-R were used to clone the first gDNA half. ABHD1 cDNA was cloned into a pCR-Blunt II-TOPO (Invitrogen) plasmid using primers cABH1_F and cABH1_3U-V_R on the previous retro-transcribed total RNA and was sequenced. Primers ABH1-CE3 and ABH1-C3U were used to clone the second cDNA half. By using an In-Fusion HD cloning kit (Clontech), the two fragments were fused in a pCR-XL-TOPO (Invitrogen) plasmid containing the *AphVII* cassette conferring resistance to hygromycin.

For subcellular protein localizations, the mCherry protein was fused either at its N-terminus or C-terminus to the above hybrid *ABHD1* coding sequence, and both driven by the *PSAD* promoter. mCherry was chosen because it emits in the orange part of the spectrum and is therefore compatible with the use of green LD-marker dye BODIPY (11) and chlorophyll autofluorescence in the red. We employed the Chlamy MoClo kit (12) and major steps are detailed below. Two different transcription units (TU) were assembled in level 1 recipient pICH47732 using the *Bsa*I enzyme: TU1, mCherry fluorescent protein cloned in-frame to ABHD1 C-terminus, and TU2, mCherry fused to ABH1 N-terminus. For both TU1 and TU2, level 0 parts used were p0-01 (*PSAD* promoter) at A1/B1 position and p0-06 (*PSAD* terminator) at B6/C1 position. ABH1 hybrid sequence was amplified by primers hABH1_B2-B4_F and hABH1_B2-B4_R to make the B2-B4 module in TU1, and primers hABH1_B5_F and hABH1_B5_R to make the B5 module in TU2. Level 0 part mCherry with RBCS2i1 +STOP at B5 position was used in TU1, and was amplified with primers mCherry_B2-B4_F and mCherry_B2-B4_R to make the B2-B4 module for TU2. Each of these TUs was cloned in level 2 at 1F position, with the *AphVII* cassette at 2F position and linker element pICH41744. The three modules were assembled into acceptor pAGM4723 using the *Bpi*I enzyme. Resulting level 2 plasmids p2-ABH1-TU1 and p2-ABH1-TU2 were digested with *Nde*I and *Bam*HI producing 6.9-kb fragments that were purified from gel and used for transformation. The chimeric *ABHD1* was cloned behind the strong promoter *PSAD* and the resulting construct was transformed by electroporation to UVM4, a strain known to enhance transgene expression due to its attenuated silencing mechanisms (13).

**Genetic transformation, screening and Confocal microscopy**

*Chlamydomonas reinhardtii* strain UVM4 generated by (13) was used for the subcellular protein localization experiment and as the host strain to generate the overexpressors. UVM4 strain was grown in TAP to about $5\times10^6$ cells $mL^{-1}$, harvested and concentrated in aliquots of 250 µL containing $7.5\times10^7$ cells, and transferred to 0.4 cm-gapped cuvettes. Linearized p2-ABH1-TU1 or p2-ABH1-TU2 (500 ng each) were added and placed on ice. Electroporation was set at 700 mV, 25 µF with no shunt resistance in a Gene Pulser (Bio-Rad). Immediately after the pulse, cells were transferred into TAP supplemented with 60 mM sucrose and were kept overnight in dim light to recover. Then cells were gently plated onto selective TAP agar supplemented with 10, 20 or 30 µg $mL^{-1}$ hygromycin under low light. Transformants were precultured in liquid TAP medium. Exponentially growing cells were screened in a TECAN Infinite M200 (Life Sciences) plate reader for mCherry fluorescence (excitation 550 nm, emission 600 nm).

To confirm mCherry signal in positive clones selected through the TECAN plate reader, they were subsequently observed under a Confocal laser scanning microscope (ZEISS LSM780) under a 63× oil immersion objective. BODIPY 505/515 (4,4-difluoro-1,3,5,7-tetramethyl-4-bora-3a,4a-diaza-s-indacene; D3921, ThermoFisher Scientific) stock at 1 mg $mL^{-1}$ in DMSO was used at a 1:100000 dilution on cell samples and incubated at room temperature for 5 min to stain neutral lipids. An argon laser line at 488 nm was used to capture fluorescence emissions of BODIPY (495-530 nm) and chlorophyll (650-680 nm), and a diode-pumped solid-state laser (DPSS) at 561 nm was used to excite mCherry (591-626 nm) in a separate track to avoid signal overlap with BODIPY emission from the 488 nm excitation. Images were obtained using Zen Black software (Carl Zeiss).

ImageJ particle analysis was used for mCherry colocalization and for the study of LD abundance and size. At least 5 images per condition of BODIPY-stained cells were taken at the same amplification, laser and fluorescence intensities and when most of cells were at their median plane (largest chlorophyll area). Cell number and areas were defined with the chlorophyll channel. LD were counted within the cell areas defined by the BODIPY channel, and diameters were calculated from LD measured areas:

$$area = \pi \left(\frac{diameter}{2}\right)^2$$

In the colocalization study, a signal prominence cutoff was set using 105 UVM4 WT cells as negative control to distinguish between autofluorescence and real mCherry signal.

**Cloning cDNA for expression in *E. coli*, purification of recombinant ABHD1 protein, refolding and activity assays**

***Gene cloning for expression in E. coli***

ABHD1 sequence was tailored for heterologous expression in *E. coli*. First, codon usage was adapted by PCR mutagenesis: rare Arg codon CGA was mutated to CGT with primers cABH1_mutR_F and cABH1_mutR_R. Second, the N-terminal (1-46 aa) was excluded since it was predicted by TMHMM software (3) as transmembranal, and amino acid residues from 47 to the last 490 aa were cloned into pLIC03 vector ($Kan^R$), which has a pET28 base adapted for ligation independent cloning (LIC). This fragment was introduced downstream of the ATG start codon of a 6xHis tag and a *tobacco etch virus* (TEV) protease-cleavage site (MGHHHHHHSSGVDLGTENLYFQSM). The construct was cloned with primers cABH1_pLIC_F and cABH1-t_pLIC_R and transformed into production strain *E. coli* ArcticExpress (DE3). To create the H356A mutant, the cDNA was altered by PCR mutagenesis: His356 codon CAC was mutated to GCC Ala with primers cABH1_mutH_F and cABH1_mutH_R (***SI Appendix*, Table S2**).

***E. coli cultures and partial purification of ABHD1 in the soluble fraction***

The recombinant protein was produced in ArcticExpress (DE3) *E. coli* cells cultured in TB medium at 37°C up to an OD of 1. At this stage, cultures were induced with 500 µM IPTG and the temperature was lowered to 17°C. Cells were grown for an additional 24 h. The cells were then harvested by centrifugation at 4000 *g* for 30 min and the pellet was frozen. For purification, cell pellet was resuspended in lysis buffer (10 mL of lysis buffer for one liter of cells at OD=1) containing 300 mM NaCl, 50 mM Tris pH 8.0, 10 mM imidazole, 5% (w/v) glycerol, 0.25 mg $mL^{-1}$ lysozyme, 20 mM $MgSO_4$, 10 µg $mL^{-1}$ DNase, and antiproteases (SIGMAFAST™ Protease Inhibitor Tablets). After resuspension, cells were lysed by sonication (sonicator probe at a frequency of 20 kHz, four 45-sec cycles separated by a one-minute delay) and centrifuged during 30 min at 12000 *g*. Supernatant was collected and enzyme was purified by fast protein liquid chromatography (FPLC) (ÄKTA Cytiva) using a nickel-NTA column (HisTrapTM HP). Protein was eluted by a step gradient using 50% (v/v) of a second buffer containing 300 mM NaCl, 50 mM Tris pH 8.0, 500 mM imidazole 5% (w/v) glycerol. The protein was concentrated and buffer was changed by repeating concentration step in a buffer containing 150 mM NaCl, 10 mM Tris pH 8.0, 5% (w/v) glycerol using Amicon ® Ultra Centrifugal filters (30 kDa). rABDH1 protein concentration was estimated using absorbance at 280 nm (ε = 1.658 g $L^{-1}$) and purity on gel. Protein rABDH1 partially purified at a concentration of about 1.7 mg $mL^{-1}$ was flash-cooled in liquid nitrogen and stored at -80°C until use for initial optimization of activity (Figures S10, S11).

***Purification of ABHD1 from inclusion bodies***
To further purify the rABHD1 protein and also enhance the productivity, pellet from the lysis step containing the inclusion bodies was resuspended in washing buffer Tris 50 mM pH 8, NaCl 300 mM, Triton X-100 1%, urea 1 M and grinded with ULTRA-TURRAX T25 until lump disintegration. Inclusion bodies were centrifuged at 12000 *g*, 30 min at 4°C and resuspended in a second washing buffer containing Tris 50 mM pH 8, NaCl 300 mM, urea 1 M. Pellet was grinded again thoroughly and centrifuged at 12000 *g*, 30 min, 4°C. Last washing step with grinding was repeated in case pellet was not yet uniformly grey. Homogenized pellet was resuspended with denaturing buffer Tris 50 mM pH 8, NaCl 300 mM, imidazole 10 mM, guanidine hydrochloride 8 M by grinding again. Solubilized inclusion bodies were micro-filtered in a Nalgene Filtration unit (0.2 µm). Denatured rABHD1 was purified in the same way as the soluble rABHD1 except that buffers were added with 6 M urea and a Nickel Chelating Resin (G-Biosciences) was used with a contact time of 30 min. For gel electrophoresis, fractions containing guanidine hydrochloride were dialyzed against water prior to sample reduction. Eluted protein was concentrated and imidazole removed using an Amicon Ultra 30K tube. A panel of solutions was tested based on recovery of lyso-PC lipase activity (***SI Appendix*, Table S3**). Refolding solution Tris 50 mM pH 8, n-octyl beta-D-glucopyranoside (O-8001, Sigma) at 10 mM was the best working combination. Denatured rABHD1 protein was refolded by performing a slow 1:20 dilution: concentrated protein was dripped on refolding solution while mixing on ice. Refolding took place overnight on a rotating wheel at 4°C. Precipitated protein was removed by centrifugation at 20000 *g* for 10 min at 4°C. Alternatively, denatured rABHD1 was dialyzed overnight against Tris 50 mM pH 8 and used as control for the droplet-embedded vesicle experiments (non-refolded urea-purified rABDH1 was still soluble after dialysis but showed 100% loss in lipase activity).

***Generation of anti-ABHD1 antibodies and immunoblot***
To generate antibodies against Chlamydomonas ABHD1, purified protein (3.4 mg) in Tris 50 nm, NaCl 300 nm buffer pH 8 were deep-frozen and shipped to ProteoGenix SAS (France) (www.proteogenix.science) for polyclonal antibody production in rabbits. SDS-PAGE gels were blotted onto nitrocellulose membranes (BioTrace NT, Sigma-Aldrich) by Semi-Dry transfer (Bio-Rad) for 75 min at 10 V using a Tris Glycine SDS buffer. Ponceau's red was applied over the membrane to check transferred proteins and Coomassie blue to observe non-transferred proteins in gel. Membranes were blocked for 30 min with Tris-Buffered Saline solution containing 0.1% Tween 20 (TBST) supplemented with 5% (w/v) nonfat dry milk. Primary antibody anti-ABHD1 generated in this study was incubated at room temperature for 4 h at 1:1000 dilution. Rabbit anti-His tag antibody was incubated for 2 h at 1:2000. Secondary antibody anti-rabbit horseradish peroxidase (HRP)-conjugated antibody (Sigma-Aldrich) was incubated for 45 min at 1:10000 dilution. Membrane was washed three times at each step with TBST and before antigen detection. Immobilon Western Chemiluminescent HRP Substrate (EMD Millipore) peroxide solution and luminol reagent were applied to the membrane, and images were recorded in a G:BOX Chemi XL (Syngene).

***Protein electrophoresis for quantification***
Purified rABHD1 protein and BSA at different concentration was denatured by incubation at 70°C for 20 min in NuPAGE LDS sample buffer with 1 mM dithiothreitol. Proteins were separated on a 12% (w/v) acrylamide Bis-Tris gel using a MOPS buffer. After staining with ProSieveTM EX Safe Stain, quantification of total protein content in gel was done using an infrared imaging scanner measuring 700-nm fluorescence.

***Enzymatic reactions***
All assays were performed in transparent glass vials sealed using caps with septum. The lipid substrate (30 µL at 1 mg $mL^{-1}$) was evaporated in a glass vial under a $N_2$ stream and 500 µL of buffer with detergent at 1 CMC was added. The reaction mixture was vortexed, sonicated for 30 s and 5 µL of purified enzyme at 1.7 mg $mL^{-1}$ (33.6 µM) was added and the mixture incubated at 25ºC for 1 – 24 h under mild agitation. Lyso-PC was used as substrate for most experiments. Lyso-DGTS, lyso-PE, lyso-PG and MAG were also tested as substrates. All lipids except lyso-DGTS were obtained from Sigma or Larodan.
The effect of pH (6 – 8.5) and sodium chloride concentration (0 – 500 mM) on enzyme activity was first evaluated using Teorell Stenhagen universal buffer and led to the choice of pH 7.5 and 100 mM NaCl. A range of buffers (Bis Tris Propane, 3-(N-morpholino)propanesulfonic acid (MOPS), phosphate, Tris-HCl and Teorell), as well as detergents at 1 CMC (Triton-X100, Lauryldimethylamine oxide (LDAO), Sodium taurodeoxycholate, Octyl β-D-glucopyranoside and 3-((3-cholamidopropyl) dimethylammonio)-1-propanesulfonate (CHAPS)) and cations at 1 mM ($ZnCl_2$, $CaCl_2$ and $MgCl_2$) were then tested. These results are presented in ***SI Appendix*, Fig. S10**. Finally, the conditions chosen to further characterize rABDH1 activity (Fig. 4 and Fig. S11) are the following: Teorell Stenhagen universal buffer (33 mM citric acid monohydrate, 33 mM phosphoric acid, 16.7 mM boric acid, 50 mM NaOH, pH 7.5 adjusted with 1 N HCl) with 100 mM NaCl and Triton-X100 at 1 CMC.

***Substrate and product determination by UPLC-MS/MS***
Reactions were stopped by lipid extraction (see details in 'lipid extraction' section). To protonate fatty acids, 1 µL of formic acid was added to the aqueous phase. Standards MAG 18:1 (0.5 µg) and oleic acid (0.5 µg) were also added to evaluate extraction efficiency. The organic phase containing lipids was dried under a flow of $N_2$, and dried lipids were re-dissolved in 100 µL of acetonitrile/isopropanol/ammonium formate 10 mM (65:30:5, v/v/v) for UPLC-MS/MS analysis.

***Isolation of lyso-DGTS from total Chlamydomonas lipid extracts for being used as substrate for enzymatic activity assays***
Lyso-DGTS (16:0) was purified from a total lipid extract from *Chlamydomonas* using UPLC ultimate RS 3000 (Thermo Fisher, Waltham, MA, USA) connected to a quadrupole-time-of-flight (QTOF) 5600 mass spectrometer (AB Sciex, Framingham, MA, USA). Lipid extracts were separated on a KinetexTM C8 2.1 × 150 mm 1.7 µm column (Phenomenex, Torrance, CA, USA). Two solvent mixtures, acetonitrile-water (60:40, v/v) and isopropanol-acetonitrile (90:10, v/v), both containing 10 mM ammonium formate at pH 3.8, were used as eluent A and B respectively. The elution was performed with a gradient of 32 min; eluent B was increased from 27 to 97% in 20 min then maintained for 5 min, solvent B was decreased to 27% and then maintained for another 7 min for column re-equilibration. The flow rate was 0.3 mL $min^{-1}$ and the column oven temperature was maintained at 45°C. First, identification of lyso-DGTS 16:0 was based on mass accuracy peaks from the MS survey scan and fragment ions from MS/MS scan. After, several injections were run and the lyso-DGTS 16:0 was collected between the retention time 2.2 and 2.8 min. The collected fractions were pooled and evaporated to dryness under nitrogen gas. The lyso-DGTS 16:0 was redissolved with a mixture of chloroform/methanol 2:1 (v/v).

***Preparation of lyso-DGTS from commercial DGTS***
1-palmitoyl-sn-glycero-3-O-4'-(*N,N,N*-trimethyl)-homoserine was obtained by partial hydrolysis of 1,2-dipalmitoyl-sn-glycero-3-O-4'-(*N,N,N*-trimethyl)-homoserine (1607456-57-4 Avanti Polar Lipids). DGTS (100 µL at a concentration of 1 mg $mL^{-1}$ in chloroform/methanol/water 2/1/0.3, v/v/v) was added to 900 µL of water containing sodium hydroxide (1 mM) and stirred during 20 min. The resulting mixture was loaded on SPE C18 cartridge and 2 mL of a solution of acetonitrile/isopropanol/water (60/35/5, v/v/v) was used to elute palmitoyl-lyso-DGTS. The resulting solutions of palmitoyl-lyso-DGTS were evaporated and resuspended in 1 mL of acetonitrile/isopropanol/water (60/35/5, v/v/v) for enzymatic assays.

**Lipid extraction and analysis**
Lipids from *Chlamydomonas* cells, isolated LDs or after enzymatic assays were extracted using a protocol based on isopropanol and methyl tert-butyl ether (MTBE) (14). Samples were collected in glass tubes by centrifugation (3200 *g*, 5 min, 4ºC), and were homogenized by vortexing and sonication in isopropanol containing 0.01% (w/v) butylated hydroxytoluene (BHT), important to protect lipids from oxidation, and kept at -20°C until further analysis. Phase separation was obtained by adding MTBE and water in a ratio of isopropanol/MTBE/water (1:3:1, v/v/v). Mixture was vortexed well and centrifuged at 4°C for 2 min at 3200 *g*. Organic upper phase was collected to a clean tube using a Pasteur pipette and one additional vol of MTBE was added to extract remaining lipids. The combined organic phases were evaporated under a gentle stream of $N_2$ and re-dissolved into a solvent mixture.

For HP-TLC, samples were re-dissolved in chloroform/methanol (2:1, v/v). Lipid classes were first separated and then quantified using HPTLC as described in (15). Total fatty acid amount in cells was analyzed by transmethylation of the lipid extract. The produced fatty acid methyl esters (FAMEs) were separated by Gas Chromatography coupled to Mass Spectrometry (GC-MS) for identification and to Flame Ionization Detector (GC-FID) for quantification as detailed in Légeret et al. (14). For LC-MS/MS, lipid extracts were dissolved in an acetonitrile/isopropanol/ammonium formate 10 mM (65:30:5, v/v/v) mixture. TAG 17:0/17:0/17:0, PE 17:0/17:0, MGDG 18:0/18:0 (0.8 µg each), MAG 18:1 or lyso-PC 17:0 (0.5 µg each), and/or oleic acid (0.05 µg) (Sigma, Larodan) were used as extraction standards as in (14).

**In vitro model droplet studies**
GUVs and incorporation of TAG emulsion are done as described in Chorlay *et al*. STAR Protocols, 2020 (16). GUVs are produced by electroformation on ITO (Indium Tin Oxide) plates with the following lipid composition: DOPC/LysoPC/Rhodamine-PE 79.5/20/0.5% (w/w) and DOPC/Rho-PE 99.5/0.5% (w/w). Formation is done in a sucrose solution with osmolarity around 270 mOsm. Buffer used for observation is a HEPES-potassium acetate-magnesium chloride buffer (HKM) composed of: 50 mM HEPES, 120 mM potassium acetate and 1 mM $MgCl_2$, with pH adjusted to 7.4 and osmolarity around 270 mOsm. Emulsion of NBD-TAG is prepared by adding 3 µL of NBD-TAG in 70 µL of buffer HKM and consecutively vortexing and sonicating the solution until it turns milky. 50 µL of the emulsion is placed on 50 µL of the GUVs solution and gentle mixing is allowed on a spinning wheel for 10 min. HKM buffer (180 µL) is deposited on a glass coverslip (Menzel-Gläser) and 20 µL of the GUVs solution is gently added. GUVs sediment and membrane breaks on the glass coverslip within 5 min. The sample is observed with a confocal Zeiss microscope, a GUV is selected and a first image at the membrane plane is taken as a reference (corresponds to time t = -5 min in **Fig. 5**). The purified recombinant ABHD1 is added at t = 0, 1 or 12 µL of solution at 1.7 mg/mL, and homogenised by pipetting. Time-lapse acquisition is started 1 min after protein addition and runs for 45 min with one image per minute, at the membrane plane. Analysis is performed with ImageJ. The surface of the droplet is measured from the NBD-TAG signal at the membrane plane according to time. Results are normalized by the initial size of the droplet at t = -5 min.

**LD isolation and purification**
To isolate LDs, we used a protocol modified based on (17). Briefly, N-starved cells of *Chlamydomonas* ($2\text{-}4\times10^9$) were pelleted at 1000 *g* for 10 min at 4°C (Avanti J-26 XP, rotor JL-8000, Beckman Coulter). The cell pellets were resuspended in 13 mL Hepes-KOH buffer (25 mM, pH 7.5) (used throughout all the purification steps) containing 0.6 M sucrose and 83 µL of Plant protease inhibitor cocktail [4-(2-aminoethyl) benzenesulfonyl fluoride (AEBSF), bestatin, pepstatin A, E-64, leupeptin, and 1,10-phenanthroline] (Sigma). In order to break the cells, samples were passed twice through a French press (Thermo Electron Aminco) at 10 MPa. In an ultracentrifuge tube with 10 mL of buffer containing 0.4 M sucrose, the cellular lysate was slowly syringed at the bottom in order to obtain two unmixed layers. LDs were separated in this sucrose gradient by centrifuging at 10000 *g* for 30 min at 4°C (Optima L-80 XP Ultracentrifuge rotor SW 32 Ti, Beckman Coulter) with a swing-out rotor. With the aid of a cut 1-mL filter tip, floating fraction containing LDs (3 mL) were collected and mixed with 9 mL of buffer containing 0.2 M sucrose and 0.1% Tween-20. In another ultracentrifuge tube with 12 mL of buffer, detergent-washed LD fraction was syringed at the bottom to obtain again two unmixed layers. After centrifuging at 10000 *g* for 30 min at 4°C (Rotor SW 32 Ti), 1 mL of upper lipid pad was collected again and mixed with 4 mL of buffer containing 0.6 M sucrose and 2 M NaCl. The fraction containing LDs were layered

in a clean ultracentrifuge tube under 5 mL buffer containing 0.2 M sucrose and 2 M NaCl with a syringe. Tubes were centrifuged at 10000 *g* for 30 min at 4°C (Rotor SW 32.1 Ti) and 1 mL of lipid pad layered on top was collected, mixed with 5 mL of urea 9 M, and incubated at RT for 10 min with gentle agitation. Then the solution was syringed at the bottom of a clean ultracentrifuge tube to layer 5 mL of buffer on the top and samples were centrifuged at 10000 *g* for 30 min at 4°C (Rotor SW 32.1 Ti). About 1 mL of washed LDs on the surface were collected and stored at -80°C for further analysis.

Polar and neutral lipid fractions were separated by solid-phase extraction in an $NH_2$ aminopropyl-modified silica Chromabond 3 mL/200 ng (Macherey-Nagel). Column was equilibrated with 5 column vol of hexane, then loaded with LD sample in chloroform/methanol (2:1, v/v) mixed with hexane (hexane/sample 20:1, v/v). Column was washed with 1 vol of hexane and 1 vol of chloroform. Flowthrough and washed fractions were pooled and dried under $N_2$ to analyze neutral lipids. Polar lipids were eluted with 2 vol of methanol and 50 mM ammonium formate. The latter fraction was re-extracted with the modified isopropanol and MTBE method before polar lipid analysis.

**Protein analyses**

***Protein extraction***

To precipitate proteins from isolated LDs, samples were first vortexed with SDS to a final concentration of 1% (v/v) at RT. Then 9 vol of pure cold acetone at -20°C was added to 1 vol of sample and samples were stored overnight at -20°C. Protein precipitates were centrifuged at 18000 *g* for 10 min at 4°C, supernatant removed and remaining acetone dried under the hood. For electrophoresis, proteins were directly resuspended in NuPAGE denaturing buffer with reducing agent following the NuPAGE protocol (Thermo Fisher): samples were loaded in 4-12% Bis-Tris Mini Gel and run with MES SDS running buffer.

***Silver nitrate staining***

Gel was fixed in a 20 mL ethanol, 5 mL acetic acid, 25 mL water fixating solution for 30 min and then transferred into an incubation solution (15 mL ethanol, 3.4 g sodium acetate, 260 µL glutaraldehyde 25% (w/v) and 0.1 g sodium thiosulfate $5H_2O$ in 50 ml water solution) for 30 min. Gel was washed three times for 5 min with water before staining with silver solution (50 mg silver nitrate and 15 µL formaldehyde in 50 mL water solution). After 30 min incubation, staining was performed with a development solution (2.5 g sodium carbonate and 13 µL formaldehyde in 100 mL water solution) applied in two baths. The first bath was discarded when solution turned grey, and the second until spots intensified for several minutes, then the gel was incubated in a stopping solution (0.73 g EDTA-$Na_2$ in 50 mL water solution) for 10 min and was washed again in water.

***MS-based quantitative proteomic analysis of LDs***

Three biological replicates from two independent LD isolations were analyzed for CC4533 (WT), *abhd1-1* and *abhd1-2* LDs. The nine samples were normalized by total fatty acid amount and further corrected using gel scanning and total band quantitation to load the same amount of protein in a silver-stained gel. The proteins, solubilized in Laemmli buffer, were stacked in the top of a 4-12% NuPAGE gel (Invitrogen). After staining with R-250 Coomassie Blue (Biorad), the proteins were digested in-gel using trypsin (modified, sequencing purity, Promega), as previously described (18). The resulting peptides were analyzed by online nanoliquid chromatography coupled to MS/MS (Ultimate 3000 RSLCnano and Q-Exactive HF, Thermo Fisher Scientific) using a 140 min gradient. For this purpose, the peptides were sampled on a precolumn (300 µm x 5 mm PepMap C18, Thermo Scientific) and separated in a 75 µm x 250 mm C18 column (Reprosil-Pur 120 C18-AQ, 1.9 µm, Dr. Maisch). The MS and MS/MS data were acquired by Xcalibur (Thermo Fisher Scientific). Results are presented in **Dataset S1**.

Peptides and proteins were identified by Mascot (version 2.8.0, Matrix Science) through concomitant searches against the Chlre5_6 database (downloaded from JGI Genome Portal (19) and a homemade database containing the sequences of classical contaminant proteins found in proteomic analyses (keratins, trypsin, etc.). Trypsin/P was chosen as the enzyme and two missed cleavages were allowed. Precursor and fragment mass error tolerances were set at respectively at 10 and 20 ppm. Peptide modifications allowed during the search were: Carbamidomethyl (C, fixed), Acetyl (Protein N-term, variable) and Oxidation (M, variable). The Proline software (20) was used for the compilation, grouping, and filtering of the results (conservation of rank 1 peptides, peptide length ≥ 6 amino acids, false discovery rate of peptide-spectrum-match identifications < 1% (21), and a minimum of 1 specific peptide per identified protein group. Proline was then used to perform the MS1 quantification, based on razor and specific peptides, of the identified protein groups.

Statistical analysis was then performed using the ProStaR software (22). Proteins identified in the contaminant database, proteins identified by MS/MS in less than two replicates of one condition, and proteins detected in less than three replicates of one condition were discarded. After log2 transformation, abundance values were normalized using variance stabilizing normalization, before missing value imputation (slsa algorithm for partially observed values in the condition and DetQuantile algorithm for totally absent values in the condition). Statistical testing was then conducted using limma, whereby differentially expressed proteins were sorted out using a log2(fold change) cut-off of 1 and a log10(p-value) cut-off of 1.4, leading to a FDR inferior to 5% according to the Pounds estimator. Proteins found differentially abundant but identified by MS/MS in less than two replicates, and detected in less than three replicates, in the condition in which they were found to be more abundant were invalidated (p-value = 1).

The MS proteomics data have been deposited to the ProteomeXchange Consortium via the PRIDE (23) partner repository with the dataset identifier (PXD036778).

***Analysis of lipid flux in N deprived Chlamydomonas using [14C]acetate***

Pulse-chase labeling experiments were completed on N deprived WT (CC4533 and UVM4) as well as *abhd1-1 abhd1-2* Chlamydomonas following the protocol described in (24). Briefly, Chlamydomonas cells grown to mid-log phase in TAP medium were spun down at 1500 g for 3 min and transferred to 50 mL TAP without N (described above) at an optical density of 0.1 at 750 nm, and grown 12 h under standard conditions to induce N deprivation. Next, Chlamydomonas were spun down again and transferred to a sterile 8 mL culture tube containing 5 mL TAP without N medium containing only 6 mM unlabeled acetate and an additional 5 µCi/ml [14C]acetate (specific activity 54.2 mCi/mmol; American Radiolabeled Chemicals Inc, St. Louis, Mo) and labeled for 3 h. During labeling 0.5 mL of suspended cells were collected at 20, 60, and 180 min into 8 mL glass vials with PTFE lined caps. Following collection of the 180 min time point, Chlamydomonas were spun down at 1500 g for 3 min and washed with 5 mL unlabeled media three times. Then, cells were transferred into 50 mL flasks containing 35 mL TAP without N and grown under standard conditions (see above). Samples comprised of 5 mL of cell suspension were collected in 8 mL glass vials with PTFE lined caps during the chase at 2, 5, 9, 21, 45 and 69 post uptake.

Following sample collection, the optical density (750 nm) of cells was measured in 5 mL total volume TAP without N. Cells were then concentrated by centrifuging at 3000 g for 3 min and the supernatant pipetted off. Lipid extracted was achieved by the addition of 3 ml chloroform/methanol/formic acid (2:1:0.1, v/v) and vortexing. Next an aqueous wash containing 0.75 mL 0.2 M $H_3PO_4$ and 0.5 mL 1 M KCl was added, samples vortexed, and centrifuged at 3000 g for 5 min to force phase separation. The upper aqueous phase was removed, and the aqueous wash was repeated twice to remove excess [14C]acetate. The lower organic phase was transferred to a new 8 mL glass vial and the remaining lipids were back extracted by adding an additional 2 mL of chloroform and repeating centrifugation and transfer steps. Finally, samples were blown down under N2 stream and re-eluted in 0.5 ml toluene + 0.005% BHT to protect samples from oxidation. Measurement of [14C]labeled lipids was achieved using a 1260 Infinity HPLC system (Agilent, Santa Clara, CA) coupled with β-Ram 6 flow-cell radiation counting module as described in (25). The identity of radioactive lipids were determined based off of HPLC retention time of unlabeled lipid class standards (from Avanti Polar Lipids, or Sigma) and measured by an Evaporative Light Scatter Detector (ELSD) as in (26).

**Yeast expression analysis**

***Plasmid construction***

For expression vector construction in yeast, the *ABDH1* sequence was optimized as the codon usage of *Saccharomyces cerevisiae* and synthesized with *Kpn*I at N-terminus and *Eco*RI at C-terminus. The products were then subcloned into pYES2 vector (Invitrogen) to form Cre12-pYES2 for expression in *S. cerevisiae*. As a positive control in yeast expression assays, the yeast diacylglycerol acyltransferase 1-coding gene (*ScDGA1*) was cloned, in a manner similar to *ABHD1*, to form pXJ412.

***Yeast strains and cell culture***

*S. cerevisiae* strain H1246 (relevant genotype: *MATαare*1-Δ::*HIS3are2*-Δ:: *LEU2 dga1*-Δ::*KanMX4lro1*-Δ::*TRP1ADE2*) containing knockouts of *DGA1*, *LRO1*, *ARE1* and *ARE2* (27) was kindly provided by S. Stymne (Scandinavian Biotechnology Research, Alnarp, Sweden). It is a neutral lipid-deficient quadruple knockout mutant for *ScDGA1*, lecithin cholesterol acyl transferase related open reading frame1 *LRO1*, acyl-coenzyme A: cholesterol acyl transferase-related enzyme 1-coding gene *ARE1* and acyl-coenzyme A: cholesterol acyl transferase-related enzyme 2-coding gene *ARE2*. Yeast cells were maintained on YPD plates (1% yeast extract [w/v], 2% peptone [w/v], and 2% glucose [w/v]) solidified with 2% agar (w/v). Cells were transformed using the lithium acetate procedure (28) and transformants were selected by growth on synthetic glucose medium (2% glucose [w/v] and 0.67% yeast nitrogen base without amino acids [w/v]) containing appropriate auxotrophic supplements (Clontech). Single yeast colonies were inoculated into liquid synthetic glucose medium and cultured overnight at 30°C and 150 rpm in an orbital shaker. The $OD_{600}$ of the culture was determined, an appropriate volume of cell culture was harvested by centrifugation, and cells were resuspended in synthetic galactose medium (2% galactose [w/v], 1% raffinose [w/v], 0.67% yeast nitrogen base without amino acids [w/v], and appropriate auxotrophic supplements) at an $OD_{600}$ of 0.4. Following expression, $1.0\times10^{11}$ yeast cells at late stationary phase of growth were used for extraction of total lipids for further analysis.

***Lipid isolation and quantification***
Total lipids were extracted from dried samples using chloroform:methanol (2:1 [v/v]) with 100 mM internal control tri13:0 TAG (Sigma) and separated on a silica TLC plate using a mixture of solvents consisting of petroleum ether, ethyl ether and acetic acid (70:30:1, by volume). To quantify the amount of TAG accumulated in yeasts expressing the CrABHD1 constructs, TAG bands were scraped from the TLC plate. Fatty acid methyl esters (FAMEs) were prepared by acid-catalyzed transmethylation of the TAG bands and then analyzed by GC-MS as previously described (29). Mixed analytical standard of FAMEs (Sigma) and pentadecane (Sigma) were used as external and internal standard, respectively. The amounts of TAGs were calculated based on the results derived from GC-MS.

### Supplemental Figures

**Figure S1. Sequence and structural feature predictions for ABHD1 protein**
**A.** TMHMM predicts ABHD1 first 50 residues are in a transmembrane domain.
**B.** IUpred3 disorder tendency of each residue. High scores correspond to high probability of disorder.
**C.** Protein amino acid sequence features summarized based on predictions by TMHMM and IUPred.
**D.** 3-D structural predictions by α-fold.
**E.** Kyte-Doolittle hydropathy plot.

**Figure S2. ABHD1 phylogenetic tree construction**
MAFFT alignment of 92 Cre-ABHD1 homologues representative of major clades. GBlocks maximum length. Method PhyML (bootstrap = 100). Only shown bootstrap values >50 for Cre-ABHD1 branch in unrooted tree. Bootstrap values are below 50 at the branches, that is, in a low percentage of generated trees (less than 50 out of 100 trees), these branches are rooted as displayed below. This happens when sequence is not conserved and they branch together by chance. In other words, we cannot conclude which group is farthest in evolutionary terms: Cyanobacteria, Archaea, Halobacteria or Animals (Opisthokonta). It is mostly random which is branched closer to Cre-ABHD1. What is not random is the evolutionary distance represented by the length of the traits. Branching is only reliable in the Chlorophyceae and Trebouxiophyceae classes and we cannot draw conclusions on evolution outside these clades.

**Figure S3.** ABHD1 overexpression studies.
**A.** Gene structure for subcellular localization studies.
**B.** Oil content analysis for ABHD1 overexpressors under 2 days of N starvation. Data represents mean and standard deviation of three technical replicates.
The mCherry protein was fused at either the N- (upper) or C-terminus of the ABHD1 protein. The ABHD1 gene is driven by PSAD promoter and terminated using the PSAD terminator. Gene was

cloned with *AphVII* cassette conferring resistance to hygromycin. Abbreviations: i*: intron rbcS2i1, H/R prom: fused promoter HSP70A/RBCS2, Rt: Rbcs2 terminator.

**Figure S4.** Image analysis of additional ABHD1 overexpression lines.
Cells were cultured in TAP, then mCherry signals were detected to observe LD. Pseudo-colors were used: chlorophyll autofluorescence in blue, mCherry in red. Bar = 5 µm.

**Figure S5.** Co-localization analysis. Signal values along the red line, marked on the DIC channel, are graphed on the left. Cells were imaged in TAP-N 1 d (*N* = 105 for WT, 69 for mCherry, 218 for ABHD1-mCherry, and 479 for mCherry-ABHD1). For mCherry, 141 LD were imaged and 93 mCherry maxima did not colocalize. For ABHD1-mCherry, 154 mCherry signal maxima colocalized to 387 LD. For mCherry-ABHD1, 17 mCherry maxima colocalized to 983 LD.
Pseudo-colors were used: BODIPY-stained LDs in yellow, chlorophyll autofluorescence in blue, mCherry and mCherry-tagged ABHD1 in red. DIC: differential interference contrast. Bar = 10 µm.

**Figure S6.** Lipid and LD analysis of the knockout *abhd1* mutants 2 days after N starvation.
**A.** TAG content. TLC amounts normalized by cell volume to account for size changes among strains.
**B.** Whole cell polar lipid composition in % of total polar lipids measured by LC-MS. Data for A and B are means of three biological replicates with two technical replicates each. Bars indicate one standard deviation.
**C.** Number of LDs per cell. On the left: bars represent mean with one standard deviation. On the right: frequency histogram. *N* = 283 cells for CC4533, 170 for *abhd1-1* and 242 for *abhd1-2*.
**D.** LD diameter. On the left: bars represent mean with one standard deviation. On the right: frequency histogram. *N* = 969 LDs for CC4533, 730 for *abhd1-1*, and 1306 for *abhd1-2*.
Abbreviations: TAG, triacylglycerol; MGDG, monogalactosyldiacylglycerol; DGDG, digalactosyldiacylglycerol; PG, phosphatidylglycerol; SQDG, sulfoquinovosyldiacylglycerol; DGTS, diacylglyceryltrimethylhomoserine; PE, phosphatidylethanolamine; PI, phosphatidylinositol.

**Figure S7. Pulse-chase flux analysis of $^{14}$C-labeled lipids in N deprived Chlamydomonas**
**A.** [14C]acetate labeling of fatty acid synthesis and their distribution between different lipid classes in wild-type.
**B.** [14C]acetate labeling of fatty acid synthesis and their distribution between different lipid classes in *abhd1* mutant.
Chlamydomonas deprived of N for 12 h were pulsed with [14C]acetate for 3 hours and chased for an additional 72 h to measure lipid accumulation and changes in lipid metabolism during N starvation.

**Figure S8. SDS-PAGE of total proteins extracted from isolated LDs.**
Three independent isolations from each genotype were shown. Equal amounts of total protein were loaded based on fatty acid methyl ester (FAME) equivalents and further adjusted via whole-lane band quantitation on gel images.
MLDP: major lipid droplet protein.

**Figure S9.** In vitro enzyme assays with rABHD1 on total lipid extracts from *abhd1-1* mutants.
Legend descriptions are the same as for Figure 4B.
LC-MS/MS detected over 10000 m/z peaks for both the reaction with rABHD1 or without.
Data are the mean of three independent experiments and error bars refer to standard deviation.
Student's t-test: *** $p < 0.001$.

**Figure S10.** The hydrolysis of lyso-DGTS by refolded rABHD1 releases a fatty acid and a GTS moiety as identified by positive ion mode MS. FFA: free fatty acid.

**Figure S11.** Optimization of in vitro enzymatic activities of rABHD1.
**A**. Effect of buffers (pH 7.5).
**B**. Effect of cations and EDTA (1 mM).
**C**. Effect of detergents (1 mM) and glycerol (10% w/v).
Lyso-PC was used as substrate in all cases, and data are means of four independent reactions with standard deviations shown. Partially purified rABDH1 was used. FFA: free fatty acid.

**Figure S12.** Progress curve of rABHD1-catalyzed enzymatic reaction (A) and Effect of rABDH1 amount on product formation (B). Data are means of three replicates and error bars indicate one standard deviation. Reaction conditions: Teorell Stenhagen universal buffer pH 7.5, with 100 mM NaCl and Triton-X100 at 1 CMC. Lyso-PC was used as substrate. Partially purified ABDH1 was used. FFA: free fatty acid.

**Figure S13. Conserved residue predictions and characterization of enzymatically inactive forms of rABHD1.**
**A.** A close-up view of the 3D protein structures with the key active residues highlighted.
**B**. Lyso-lipase activity assays using rABDH1 and a H365A mutant of rABDH1. Both were soluble proteins partially purified (H365A could not be obtained under a soluble form after the refolding process of the protein urea-purified from inclusion bodies). LC-MS signal in arbitrary units (a.u.). FFA: Free fatty acid. Reactions were performed with 1.2 ± 0.6 µM enzyme, lyso-PC was used as a substrate and stopped at different incubation times by lipid extraction. Data are means of two replicates and error bars indicate one standard deviation.
**C**. Characterization of non-refolded urea-purified rABHD1 (100% inactive enzymatically) using the droplet-embedded vesicle system. Conditions for assays are the ones described in Figure 5.

### Table S1. Protein sequence IDs used to build the phylogenetic tree.

| ID | Protein ID | E-value | Score | Identity | Organism | Tree branch/Taxon | |
|---|---|---|---|---|---|---|---|
| **Cre-ABHD1** | A0A2K3D6X9_CHLRE | 0 | 1758 | 100.00% | Chlamydomonas reinhardtii | Viridiplantae | Chlorophyceae |
| **Tetrabaena-EH4** | A0A2J7ZVT9_9CHLO | 3.10E-110 | 880 | 57.80% | Tetrabaena socialis | Viridiplantae | Chlorophyceae |
| **Volvox-SYP2** | Vocar.0001s1388.1 | 3.30E-106 | 328.2 | 54.9 | Volvox carteri | Viridiplantae | Chlorophyceae |
| **Gonium-SYP2** | A0A150GU95_GONPE | 2.00E-102 | 818 | 53.10% | Gonium pectorale | Viridiplantae | Chlorophyceae |
| **Cre-ABHD2** | Cre01.g010550.t2.1 | 1.80E-92 | 294.3 | 58 | Chlamydomonas reinhardtii | Viridiplantae | Chlorophyceae |
| **C.eustigma** | A0A250WYT8_9CHLO | 1.00E-63 | 560 | 43.30% | Chlamydomonas eustigma | Viridiplantae | Chlorophyceae |
| **Coccomyxa-1** | Csu-60484 | 2.70E-57 | 198.7 | 44.4 | Coccomyxa subellipsoidea C-169 | Viridiplantae | Trebouxiophyceae |
| **Micractinium** | A0A2P6VKZ8_9CHLO | 2.60E-55 | 520 | 41.90% | Micractinium conductrix | Viridiplantae | Trebouxiophyceae |
| **Raphidocelis-EH** | A0A2V0NY35_9CHLO | 2.80E-55 | 509 | 45.40% | Raphidocelis subcapitata | Viridiplantae | Chlorophyceae |
| | A0A2W7BQR0_9CYAN | 2.80E-52 | 476 | 44.20% | Leptolyngbya sp. | Terrabacteria group | Cyanobacteria |
| | A0A0M9AP20_9EURY | 4.00E-51 | 470 | 45.30% | Haloarcula rubripromontorii | Euryarchaeota | Halobacteria |
| **Monoraphidium** | A0A0D2MKF3_9CHLO | 4.40E-51 | 479 | 45.00% | Monoraphidium neglectum | Viridiplantae | Chlorophyceae |
| | M0LQ76_9EURY | 2.80E-50 | 463 | 40.70% | Halococcus hamelinensis 100A6 | Euryarchaeota | Halobacteria |
| | M0KF12_9EURY | 3.00E-50 | 464 | 45.70% | Haloarcula amylolytica JCM 13557 | Euryarchaeota | Halobacteria |
| **Tetradesmus** | A0A383VNG8_TETOB | 4.40E-50 | 466 | 40.30% | Tetradesmus obliquus | Viridiplantae | Chlorophyceae |
| | A0A2T2W3E2_9CYAN | 6.30E-50 | 460 | 41.60% | filamentous cyanobacterium CCP3 | Terrabacteria group | Cyanobacteria |
| | A0A2R6HUZ7_9EURY | 1.80E-49 | 458 | 42.60% | Halobacteriales archaeon QS_4_62_28 | Euryarchaeota | Halobacteria |
| | J3JEL7_9EURY | 2.30E-49 | 457 | 44.40% | Halogranum salarium B-1 | Euryarchaeota | Halobacteria |
| **Sorangium** | A0A150SB42_SORCE | 3.60E-49 | 455 | 43.80% | Sorangium cellulosum | Proteobacteria | Deltaproteobacteria |
| | A0A1I6KLE4_9EURY | 3.70E-49 | 456 | 43.60% | Halomicrobium zhouii | Euryarchaeota | Halobacteria |
| **Gammaproteobacteria** | A0A2W4R6X2_9GAMM | 1.50E-48 | 452 | 39.60% | Candidatus Methyloumidiphilus alinensis | Proteobacteria | Gammaproteobacteria |
| **Rubrobacter** | A0A023X451_9ACTN | 3.30E-48 | 449 | 40.90% | Rubrobacter radiotolerans | Terrabacteria group | Actinobacteria |
| | K8GKR2_9CYAN | 3.80E-48 | 448 | 40.40% | Oscillatoriales cyanobacterium JSC-12 | Terrabacteria group | Cyanobacteria |
| | A0A2R6F938_9EURY | 1.60E-47 | 446 | 40.40% | Halobacteriales archaeon QH_8_64_26 | Euryarchaeota | Halobacteria |
| | A0A2S6V127_9CYAN | 4.80E-47 | 440 | 39.90% | Chroococcidiopsis sp. TS-821 | Terrabacteria group | Cyanobacteria |
| | A0A0B7B124_9EUPU | 1.00E-46 | 443 | 37.00% | Arion vulgaris | Opisthokonta | Metazoa |
| **D.maricopensis** | E8U4S3_DEIML | 1.50E-46 | 437 | 40.80% | Deinococcus maricopensis | Terrabacteria group | Deinococcus-Thermus |
| | A0A2D0HHR3_9NOSO | 3.80E-46 | 434 | 38.60% | Nostoc sp. DB3992 | Terrabacteria group | Cyanobacteria |
| | A0A2L2NK23_9NOSO | 5.00E-46 | 434 | 37.90% | Nostoc sp. 'Lobaria pulmonaria (5183)' | Terrabacteria group | Cyanobacteria |
| **C.variabilis** | E1ZRB9_CHLVA | 7.30E-46 | 431 | 40.60% | Chlorella variabilis | Viridiplantae | Trebouxiophyceae |
| **BTActinobac** | A0A3A4PDA9_9ACTN | 1.40E-45 | 431 | 38.00% | Actinobacteria bacterium | Terrabacteria group | Actinobacteria |
| | A0A151AGU6_9EURY | 1.90E-45 | 430 | 41.30% | Halalkalicoccus paucihalophilus | Euryarchaeota | Halobacteria |
| | V4BHS9_LOTGI | 2.40E-45 | 433 | 37.70% | Lottia gigantea | Opisthokonta | Metazoa |
| | A0A365TGG0_9EURY | 2.70E-45 | 429 | 41.70% | halophilic archaeon | Euryarchaeota | Halobacteria |
| | L9VYZ5_9EURY | 3.30E-45 | 430 | 40.40% | Natronorubrum sulfidifaciens | Euryarchaeota | Halobacteria |
| | K9SP24_9CYAN | 4.00E-45 | 427 | 39.90% | Pseudanabaena sp. PCC 7367 | Terrabacteria group | Cyanobacteria |
| | A0A1Q1FKJ6_9EURY | 4.60E-45 | 428 | 41.30% | Halorientalis sp. IM1011 | Euryarchaeota | Halobacteria |
| | A0A0D2X5E4_CAPO3 | 5.10E-45 | 431 | 37.20% | Capsaspora owczarzaki | Opisthokonta | |
| | A0A2T5K3A5_9PROT | 6.80E-45 | 429 | 42.10% | Nitrosospira sp. Nsp2 | Proteobacteria | |
| | A0A1N6XEQ0_9EURY | 8.70E-45 | 426 | 40.40% | Haloterrigena daqingensis | Euryarchaeota | Halobacteria |
| **Chlorella-1** | ChlNC64A_1\|139988 | 1.18E-44 | 550 | 43.9 | Chlorella sp. NC64A | Viridiplantae | Trebouxiophyceae |
| | A0A3N5F0Z9_9BURK | 1.70E-44 | 424 | 37.30% | Burkholderiales bacterium | Proteobacteria | |
| **Branchiostoma** | C3Y9Y3_BRAFL | 3.10E-44 | 424 | 36.90% | Branchiostoma floridae | Opisthokonta | Metazoa |
| **Auxenochlorella-EH4** | A0A087SJ87_AUXPR | 3.70E-44 | 419 | 35.90% | Auxenochlorella protothecoides | Viridiplantae | Trebouxiophyceae |

| | | | | | | | |
|---|---|---|---|---|---|---|---|
| **Nitrosovibrio** | A0A1H7JVA5_9PROT | 4.50E-44 | 420 | 42.50% | Nitrosovibrio tenuis | Proteobacteria | Betaproteobacteria |
| | A0A1W0X919_HYPDU | 6.00E-44 | 425 | 36.60% | Hypsibius dujardini | Opisthokonta | Metazoa |
| | A0A1S3HRG0_LINUN | 6.60E-44 | 424 | 35.60% | Lingula unguis | Opisthokonta | Metazoa |
| | A0A2T1GRV9_9CYAN | 1.10E-43 | 417 | 40.70% | Chroococcidiopsis cubana | Terrabacteria group | Cyanobacteria |
| | Q3MFC7_ANAVT | 1.60E-43 | 416 | 38.30% | Anabaena variabilis | Terrabacteria group | Cyanobacteria |
| **Acidobacteria** | A0A3E0PFU6_9BACT | 1.70E-43 | 416 | 39.10% | Acidobacteria bacterium | Acidobacteria | |
| **D.deserti** | C1CVI6_DEIDV | 1.90E-43 | 415 | 42.50% | Deinococcus deserti | Terrabacteria group | Deinococcus-Thermus |
| **Rokubacteria** | A0A2V6VW68_9BACT | 2.60E-43 | 415 | 37.70% | Candidatus Rokubacteria bacterium | unclassified Bacteria | |
| | A0A1Z4SVZ0_9CYAN | 3.00E-43 | 414 | 37.30% | Calothrix sp. NIES-4105 | Terrabacteria group | Cyanobacteria |
| | A0A124P884_9BURK | 3.50E-43 | 415 | 35.60% | Burkholderia singularis | Proteobacteria | Betaproteobacteria |
| **PVC bacteria** | A0A354GTM1_9BACT | 3.70E-43 | 413 | 39.90% | Planctomycetales bacterium | PVC group | |
| **Pontibacter** | A0A1I2X361_9BACT | 3.70E-43 | 413 | 38.40% | Pontibacter chinhatensis | FCB group | |
| **Spirochaetes** | A0A1G3QLK5_9SPIR | 3.80E-43 | 415 | 34.20% | Spirochaetes bacterium RBG_16_49_21 | Spirochaetes | |
| **B-Calditricha** | A0A3M2I9W3_9BACT | 8.50E-43 | 411 | 36.80% | Calditrichaeota bacterium | Calditrichaeota | |
| | M0AKL3_9EURY | 9.50E-43 | 411 | 40.00% | Natrialba chahannaoensis | Euryarchaeota | Halobacteria |
| | A0A166U6N4_9EURY | 9.70E-43 | 411 | 40.10% | Haladaptatus sp. R4 | Euryarchaeota | Halobacteria |
| **Cesiribacter** | M7N1N3_9BACT | 1.00E-42 | 411 | 39.70% | Cesiribacter andamanensis | FCB group | |
| **Blastocatellia-2** | A0A3B9MNT1_9BACT | 1.70E-42 | 411 | 37.70% | Blastocatellia bacterium | Acidobacteria | |
| **Blastobacteria1** | A0A0B6WVW5_9BACT | 2.60E-42 | 409 | 41.30% | Pyrinomonas methylaliphatogenes | Acidobacteria | Blastocatellia |
| **Chlorella-2** | ChlNC64A_1\|27475 | 3.10E-42 | 469 | 43.4 | Chlorella sp. NC64A | Viridiplantae | Trebouxiophyceae |
| **C.sorokiniana-EH** | A0A2P6TGQ4_CHLSO | 3.90E-42 | 414 | 35.90% | Chlorella sorokiniana | Viridiplantae | Trebouxiophyceae |
| **Betaproteobacteria-SG8** | A0A0S8BBK4_9PROT | 4.20E-42 | 407 | 39.20% | Betaproteobacteria bacterium SG8_39 | Proteobacteria | Betaproteobacteria |
| | A0A140K508_9CYAN | 4.60E-42 | 406 | 36.90% | Stanieria sp. NIES-3757 | Terrabacteria group | Cyanobacteria |
| **BP-methylo28** | A0A2U8WMK0_9RHIZ | 5.70E-42 | 406 | 40.60% | Methylobacterium sp. 17Sr1-28 | Proteobacteria | Alphaproteobacteria |
| | A0A1G3H5T5_9RHOO | 6.00E-42 | 406 | 36.00% | Rhodocyclales bacterium GWA2_65_20 | Proteobacteria | Betaproteobacteria |
| | A0A354SIP7_9DELT | 8.70E-42 | 404 | 34.60% | Syntrophaceae bacterium | Proteobacteria | Deltaproteobacteria |
| | A0A2B7H057_9EURY | 9.80E-42 | 405 | 39.60% | Natrinema sp. CBA1119 | Euryarchaeota | Halobacteria |
| | A0A2T7NZT4_POMCA | 1.30E-41 | 408 | 36.10% | Pomacea canaliculata | Opisthokonta | Metazoa |
| **Blastocatellia-1** | A0A321LK39_9BACT | 1.40E-41 | 403 | 36.70% | Blastocatellia bacterium AA13 | Acidobacteria | Blastocatellia |
| | V2JDL5_9BURK | 1.90E-41 | 403 | 36.00% | Cupriavidus sp. | Proteobacteria | Betaproteobacteria |
| **Bacterium-JGI053** | A0A285YXC2_9BACT | 2.00E-41 | 402 | 37.90% | bacterium JGI 053 | unclassified Bacteria | |
| | K9RN73_SYNP3 | 3.00E-41 | 400 | 37.90% | Synechococcus sp. | Terrabacteria group | Cyanobacteria |
| **Galdieria** | Galsul1\|1082 | 3.59E-41 | 471 | 43.4 | Galdieria sulphuraria 074W | Viridiplantae | Rhodophyta |
| | A0A210QE03_MIZYE | 3.90E-41 | 404 | 35.30% | Mizuhopecten yessoensis | Opisthokonta | Metazoa |
| **Coccomyxa-2** | Csu-60355 | 4.50E-40 | 151 | 38.7 | Coccomyxa subellipsoidea C-169 | Viridiplantae | Trebouxiophyceae |
| **Chlorella-3** | ChlNC64A_1\|10937 | 5.54E-40 | 467 | 48.3 | Chlorella sp. NC64A | Viridiplantae | Trebouxiophyceae |
| **Sphagnum** | Sphfalx0071s0022.1 | 4.30E-28 | 120.6 | 32.1 | Sphagnum fallax | Viridiplantae | Embryophyta |
| **N.oceanica** | Nanoce1779_2\|667590 | 5.23E-26 | 436 | 47.6 | Nannochloropsis oceanica | Chromista | Eustigmatophyceae |
| **N.gaditana** | Nangad1\|4198 | 2.98E-23 | 389 | 49.7 | Nannochloropsis gaditana | Chromista | Eustigmatophyceae |
| **Micromonas-RCC** | RCC-81226 | 2.90E-20 | 92.8 | 34.1 | Micromonas sp. RCC299 | Viridiplantae | Chlorophyta - Mamiellophyceae |
| **Phaeodactylum** | Phatr1\|50502 | 1.06E-17 | 329 | 41.2 | Phaeodactylum tricornutum | Chromista | Bacillariophyta |
| **Kalanchoe** | Kalax.0242s0009.1 | 1.10E-17 | 89.4 | 28.3 | Kalanchoe laxiflora | Viridiplantae | Embryophyta |
| **Selaginella** | Smo-266570 | 1.30E-17 | 89.4 | 26.4 | Selaginella moellendorffii | Viridiplantae | Embryophyta |
| **Citrus** | Ciclev10002315m | 5.70E-17 | 86.3 | 29.2 | Citrus clementina | Viridiplantae | Embryophyta |
| **Solanum** | PGSC0003DMT400063719 | 1.40E-16 | 85.1 | 41.7 | Solanum tuberosum | Viridiplantae | Embryophyta |
| **Ectocarpus-1** | Ectsil1\|27584 | 1.56E-14 | 241 | 44.7 | Ectocarpus siliculosus | Chromista | Phaeophyceae |
| **Ectocarpus-2** | Ectsil1\|27585 | 1.32E-13 | 235 | 43.2 | Ectocarpus siliculosus | Chromista | Phaeophyceae |
| **O.tauri** | Ostta4\|35866 | 6.92E-06 | 173 | 36.5 | Ostreococcus tauri | Viridiplantae | Prasinophyceae |

### Table S2. Primer sequences used in this study.

| Usage | Primer name | Sequence (5'-3') |
|---|---|---|
| **RT-PCR** | CBLP-F | GAGTCCAACTACGGCTACGCC |
| | CBLP-R | CTCGCCAATGGTGTACTTGCAC |
| | ABH1_E1_F | ATGTCGTCCACGGCATGTAATGGCG |
| | ABH1_E11_R | TCACTCACTGGCCGGCTGCGTC |
| **cDNA and In-fusion for hybrid ABHD1** | cABH1_F | ATCAAGCATATAACCTGAGGACGACACAA |
| | cABH1_3U-V_R | CCAACCTTACCAGAGTCCACGCCAGCGGCAG |
| | ABH1-GP-F | TATCCATCACACTGGCGGCCTGAAGAACGAGACAATCGCG |
| | ABH1-GE2-R | AGGAAGGAGCGGGGGTTGCGCCACAGGTCCCGCAG |
| | ABH1-CE3_F | GCAACCCCCGCTCCTTCCTC |
| | ABH1-C3U_R | CTCTAGATGCATGCTCGAGCGAGTCCACGCCAGCGGCAG |
| **Golden Gate (MoClo)** | hABH1_B2-B4_F | ttGGTCTCaCCATGTCGTCCACGGCATGTAATGGC |
| | hABH1_B2-B4_R | ttGGTCTCaCGAACCCTCACTGGCCGGCTGCGT |
| | hABH1_B5_F | ttGGTCTCaGGTTCGATGTCGTCCACGGCATGTAA |
| | hABH1_B5_R | ttGGTCTCaAAGCTCACTCACTGGCCGGCT |
| | mCherry_B2-B4_F | ttGGTCTCaCCATGGTGTCCAAGGGCGA |
| | mCherry_B2-B4_R | ttGGTCTCaCGAACCGAGCTCGTCGTGCTTGT |
| | cABH1_mutR_F | GACGCCCCCGAGCGTCTCAACGCCATC |

| | | |
|---|---|---|
| **Mutagenesis rare codon** | cABH1_mutR_R | GATGGCGTTGAGACGCTCGGGGGCGTCG |
| **Protein expression in *E. coli*** | cABH1_pLIC_F | TACTTCCAATCAATGCTGCTGCGGGACCTG |
| | cABH1-t_pLIC_R | TATCCACCTTTACTGTTATCACTCACTGGCCGGCTG |
| **Point mutagenesis** | cABH1_mutH_F | GCCTGCACCCAGGCGGAGGCCTCGGG |
| | cABH1_mutH_R | CCCGAGGCCTCCGCCTGGGTGCAGGC |

**Table S3. Conditions and results obtained for protein refolding work.**
**A.** Consumption of lyso-PC 17:0 calculated as a percentage of its refolding solution without enzyme.
**B.** Production of fatty acid 17:0 corrected by oleic acid extraction standard.

| Refolding solution | Buffer | Aggregation suppressor | Folding enhancer | Reducing agent | Detergent |
|---|---|---|---|---|---|
| **R2** | CHES | Arginine | Glycerol | DTT | - |
| **R8** | CHES | - | Glycerol | - | - |
| **R9** | Tris | Arginine | - | - | - |
| **R13** | MES | Arginine | - | - | - |
| **R14** | Tris | Arginine | - | - | SB3-10 |
| **R15** | Tris | Arginine | - | - | b-OG |
| **R16** | Tris | Arginine | PEG | - | - |
| **R17** | Tris | - | - | - | b-OG |
| **Previous test, solutions discarded (ranged from more to less active; data not shown):** | | | | | |
| **R11** | Glycine | Arginine | Glycerol | - | - |
| **R5** | Glycine | Arginine | - | DTT | - |
| **R6** | Tris | Arginine | Glycerol | - | - |
| **R1** | CHES | Arginine | - | DTT | - |
| **R4** | Tris | - | Glycerol | DTT | - |
| **R12** | Glycine | - | - | - | - |
| **R3** | Tris | - | - | DTT | - |
| **R7** | CHES | - | - | - | - |
| **R10** | Glycine | - | Glycerol | DTT | - |

Buffers at 50 mM: 4-Morpholineethanesulfonic acid (MES) pH 5.5, Tris-HCl pH 8, N-cyclohexyl-2-aminoethanesulfonic acid (CHES) pH 9, Glycine pH 10.
Arginine-HCl at 1 M, glycerol at 1 M, PEG: 0.05% polyethylene glycol 4000, DTT at 10 mM. Zwitterionic detergents at 10 mM: b-OG: n-octyl β-D-glucopyranoside, SB3-10: caprylyl sulfobetaine.

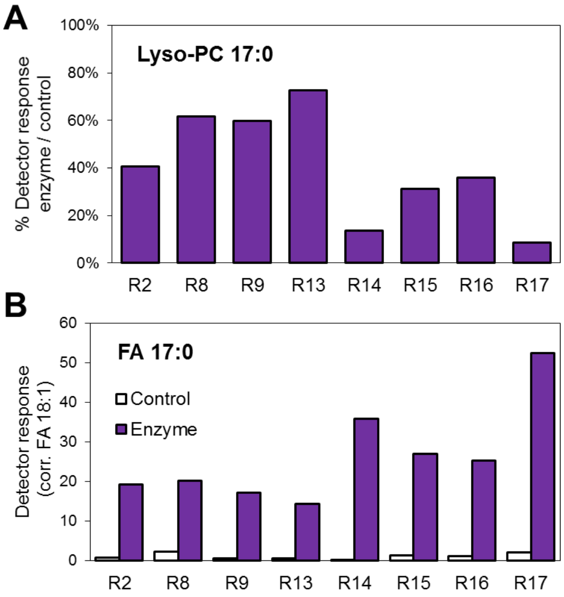


**Movie S1.** Confocal Z-stack of cells expressing ABHD1-mCherry (C-ter) fusion protein. Cells were cultured in TAP-N for 2 days. From left to right: differential interference contrast (DIC), composite image (pseudo-colors for chlorophyll autofluorescence in blue, mCherry in red, BODIPY in yellow), only mCherry, iand only BODIPY. Bar = 5 µm.

**Dataset S1.** MS-based label-free quantitative proteomic analysis of LD proteome in CC4533 WT and *abhd1* mutants.
